# Methane sink function of grassland soil microbiomes - negative effects of intensive management persist three years after land-use extensification

**DOI:** 10.1101/2025.03.31.646387

**Authors:** Nils Volles, Hauke Winter, Verena Groß, Milos Bielcik, Tim Urich, Steffen Kolb, Sven Marhan

## Abstract

Grassland soils are important methane (CH_4_) sinks through CH_4_ oxidation by methanotrophs, but intensive management with high nitrogen inputs and grazing densities reduces this potential. While long-term recovery of the CH_4_ sink after land-use change is generally established, little is known about the short-term effects of reducing land-use intensity index (LUI) through extensive management in grassland. We did not find an effect on potential CH_4_ oxidation rates (PMORs) and the abundances atmospheric CH_4_-consuming methanotrophs after three years of LUI reduction (no fertilization, no grazing, and one mowing per year) in 45 intensively managed grassland sites located in three different pedoclimatic regions of Germany. However, we observed a decline in the abundance of CH_4_ producing methanogens. Moreover, we found greater PMORs and higher abundance of Upland Soil Cluster γ (USCγ) methanotrophs on additional, historically low LUI sites. Soil bulk density decreased already after three years of LUI reduction and was even lower in historically low LUI sites. Strong correlations between the abundance canonical methanotrophs and methanogens highlight a CH_4_ filter function that was independent from LUI reduction across regions. Our study consistently shows that three years of LUI reduction are not sufficient to restore the CH_4_ sink function of temperate grasslands. However, the lower soil bulk density and the decreased abundance of methanogens indicate that LUI reduction will affect the habitat and living boundary conditions for CH_4_-cycling microorganisms in the long term.

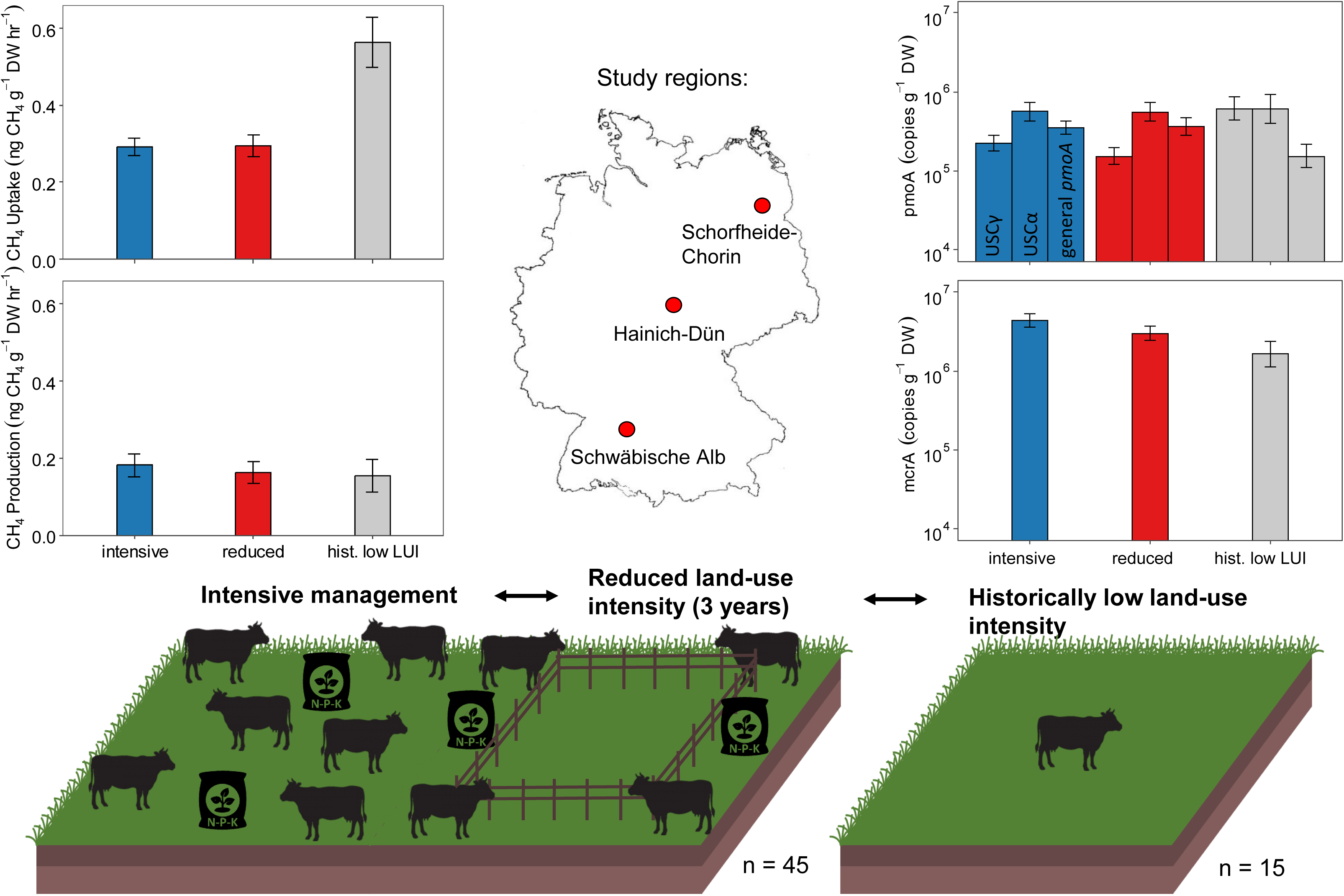

## 1. Introduction

Methane (CH_4_) concentration in the troposphere has increased 2.6-fold since the beginning of the industrial era (Thoning et al., 2024). Its contribution to radiative forcing is 30 times higher than CO_2_ calculated over 100 years, and 80 times over 20 years, respectively (Forster et al., 2021; Jackson et al., 2024). Soils contribute to the global CH_4_ cycle through two key microbial processes. Methanogenesis refers to the biological production of CH_4_ in soils and is almost exclusively mediated by methanogenic archaea (methanogens) within the Euryarchaeota. These microorganisms can be identified using the *mcrA* gene as a biomarker (Conrad, 2009; Söllinger and Urich, 2019). CH_4_ oxidation is carried out by methanotrophic bacteria (methanotrophs), which consume CH_4_ in well-aerated soils, such as those in grassland ecosystems. This process constitutes the most important terrestrial sink for atmospheric CH_4_. The particulate methane monooxygenase (pMMO) is the key enzyme in this process and the *pmoA* gene encoding its alpha subunit is used as a functional marker for methanotrophs (Knief, 2015; Baani and Liesack, 2008).

Methanotrophs in soil can function as biological CH_4_ filters by preventing the emission of a considerable proportion of CH_4_ formed by methanogens in anoxic soil compartments to the atmosphere. While these methanotrophs require high (> 600 ppm) CH_4_ concentrations to thrive, other specialized taxa such as Upland soil cluster (USC) α and γ are well adapted to atmospheric and sub-atmospheric levels of CH_4_ and thus, mediate the sink activity for atmospheric CH_4_ of soils (Täumer et al., 2021; Kolb et al., 2005). The interplay within the CH_4_-cycling microbiome therefore determines whether a soil acts as source or sink for atmospheric CH_4_. Soil environmental factors, such as pH, oxygen availability, water table level, temperature, and microbial growth substrate control the habitat boundary conditions of CH_4_-cycling microorganisms in soils (Bodelier 2011, Täumer et al. 2021).

Besides these environmental factors, land-use intensity also influences the CH_4_ sink function of grassland soils. Especially, nitrogen fertilization can reduce CH_4_ uptake of soils as proven on local (Imer et al., 2013; Mosier et al., 1991) and on global scales (Liu and Greaver, 2009). Mechanisms, such as competitive inhibition of MMO through NH_4_ ^+^, toxicity of NO ^-^ or osmotic stress can decrease abundance and activity of methanotrophs. High land-use intensity and nitrogen fertilization as its component reduce the potential CH_4_ oxidation rates (PMORs) of grasslands by 40%, as shown for 150 grassland sites in Germany (Täumer et al., 2021). Bulk density hereby constitutes a major soil characteristic modulating CH_4_ oxidation potential. Another major component of land-use intensity in grasslands is grazing. Intensive grazing reduces the atmospheric CH_4_ uptake by 24 – 31% (Pan et al., 2021; Chen et al., 2010; Abell et al., 2009).

While the negative effects of intensification are well characterized, little is known about the recovery of the CH_4_-cycling microbiome through extensification measures in grasslands. Recovery of methanotrophs populations and their activity has been studied along secondary successional gradients (Levine et al., 2011), after conversion of land-use (Ho et al., 2022; Borken et al., 2003; Dörr et al., 2010; Wu et al., 2020), and following regenerative practices in various cropping systems (Lim et al., 2024). These studies consistently report long-term CH_4_ sink recovery, occurring over decades to a century. In a few grassland ecosystems, CH_4_ surface fluxes were measured after grazing exclosure on single sites and showed increased atmospheric CH_4_ uptake after 33, and 8 years (Pan et al., 2021; Chen et al., 2010) while conflicting results were found after 4 and 6 years, respectively (Wang et al., 2023c; Wei et al., 2012; Thomas et al., 2017). Among surface fluxes, soil incubations from 6 meadow sites showed no effect after 4 years of grazing exclosure (Liu et al., 2022). Considering these studies, a systematic understanding of the short to medium term CH_4_ sink recovery from intensive management and its underlying mechanisms is still lacking.

The aim of our study was to investigate the extent to which a short-term reduction in land use intensity changes the CH_4_ source and sink function of temperate grassland soils and whether this is related to changes in the abundances and interactions of methanotrophic and methanogenic microorganisms. We studied top-and subsoil samples from 45 intensively managed grassland sites located in three pedoclimatic regions in Germany with widely varying soil properties. At each site, land-use intensity was experimentally reduced on one subplot by omitting fertilization and grazing for three years and by reducing the mowing frequency to one event per year. As a baseline for the maximum CH_4_-sink potential of grasslands, we additionally investigated 15 grassland sites with historically low land-use intensity. We quantified both CH_4_ oxidation and formation potential, the abundance of different functional groups of methanotrophs, the abundance of methanogens, and physico-chemical soil characteristics.

## 2. Materials and methods

### 2.1 Experimental design

The experiment was conducted within the framework of the Biodiversity Exploratories (https://www.biodiversity-exploratories.de; Fischer et al., 2010) with grasslands in three regions in Germany: Schorfheide-Chorin (SCH), Hainich-Dün (HAI) and Schwäbische Alb (ALB). The regions cover a broad spectrum of soil properties, altitude and climatic conditions, making them representative for a large part of Central Europe. Grasslands are used as meadows (mown at least once, if grazed then only briefly by sheep), pastures (only grazed, not mown yearly) or mixed pastures (mowing and grazing in the same year). Sampling was conducted as part of the reduced land-use intensity experiment (REX) established before the start of the vegetation period in 2020 (Andraczek et al., 2023). Here, a controlled experiment was set up at 15 grassland sites per region (total n = 45). Each experimental site consisted of an intensively managed control plot (50 × 50 m) and a reduced land-use intensity index (LUI) plot (30 × 30 m) in its close proximity. At the reduced LUI plot, no fertilization took place, grazing was excluded, and mowing was reduced to one event per year. As a baseline for the maximum CH_4_ oxidation potential, five additional grassland sites per region (total n = 15) with historically low LUI were sampled as well. LUI was calculated as regional mean of grassland management for the regions ALB, HAI, SCH for the years of 2011 to 2021 according to Blüthgen et al. (2012), based on information from the land owners on mowing, grazing and fertilization (Vogt et al., 2019) using the LUI calculation tool (Ostrowski et al., 2020) implemented in BExIS (http://doi.org/10.17616/R32P9Q). Historically low LUI plots were defined as the grassland sites with the lowest LUI in each region. At least since 2006 and likely even longer, they received no fertilization and were not grazed more than once a year.

### 2.2 Soil sampling and soil properties

In May 2023, four soil cores were collected on each grassland plot using a Split Tube (48 mm; Eijkelkamp) from topsoil (0–10 cm) and subsoil (20-30 cm). Aboveground plant material was removed before soil sampling. The four soil cores per plot were pooled and homogenized for each depth and immediately transported to the field lab in cooling bags. The soil was sieved (< 2 mm) and the mass of stones and roots in each of the pooled samples was determined. Subsequently, the sieved soil was split in subsamples for gas measurements kept at 4 °C and samples for DNA extraction and determination of soil properties kept at -20°C. Gravimetric soil water content (SWC) was determined in duplicate by drying 5 g of soil at 60°C for 72 hours. Soil pH was determined with a glass electrode (pH meter 538 and pH glass electrode SenTix 61; WTW) using 10 g of air-dried soil mixed with 25 mL of 0.01 M CaCl solution. Microbial biomass carbon (C_mic_) content was determined by chloroform-fumigation-extraction (CFE) according to Vance et al. (1987). In brief, 5 g of the field moist soil were fumigated with ethanol-free chloroform for 24 h in a desiccator and extracted in 40 ml 0.5 M K_2_SO_4_ on a horizontal shaker at 250 rpm for 30 min and centrifuged at 4400 g for 30 min. C and N concentrations in the supernatants were measured on a total organic carbon and nitrogen analyzer (multi-N/C 2100S, Analytic Jena AG). Another 5 g were directly extracted as described above but without the fumigation step (non-fumigated sample). Extracts from non-fumigated samples were used for determining extractable organic carbon (EOC), total nitrogen (ETN) and concentrations of ammonium (NH_4_ ^+^) and nitrate (NO_3_ ^−^). C_mic_ was calculated from the differences in concentrations between fumigated and non-fumigated samples using the kEC factor of 0.45 (Joergensen, 1996). The concentrations of NH_4_ ^+^-N and NO_3_ ^−^-N were measured with a continuous flow analyzer (Bran + Luebbe Autoanalyzer 3, SEAL Analytical, Hamburg, Germany) Data on soil physico-chemical parameters is stored in BExIS (http://doi.org/10.17616/R32P9Q; accession number 32036; Volles et al., 2025a).

### 2.3 Gas flux measurements of soil samples

To measure gas net fluxes of the soil samples under different conditions, 50 g for mineral soils and 70 g for organic soils were weighed into 120 ml (53 mm diameter) polypropylene plastic beakers in duplicate. Subsequently, the topsoil was re-compacted to its original bulk density (BD). The subsoil was re-compacted to a BD of 1 g cm^3^. As 20-40% of the maximum water holding capacity has been previously shown to be an optimal range for CH_4_ oxidation (Gulledge and Schimel, 1998), soil water content was adjusted to field capacity (pF 2.5) by adding deionized water or gently air-drying at 21°C. If the soil water content was different between the paired samples of intensive and reduced LUI plots, the water content was rebalanced by adding the deionized water to the drier sample. Data on soil water tension is stored in BExIS (http://doi.org/10.17616/R32P9Q; accession number 26207; Boeddinghaus et al., 2022).

To determine the activity of high-affinity methanotrophs, potential methane oxidation rate (PMOR) was measured at atmospheric CH_4_ concentrations (1.8 ppmv). For acclimation, plastic beakers were kept at 21 °C in the dark for four days before they were transferred into glass jars (500 ml; Weck Gläser; J. Weck GmbH u. Co. KG). The jars were closed with gastight lids which enabled gas sampling through three-way stopcock valves (Figure S1). 50 ml of ambient air was added at the start of the incubation to slightly over-pressurize the microcosms. Immediately, and after 2, 4, and 6 hours, 12 ml of headspace gas sample were transferred into pre-evacuated exetainers (5.9 ml; Labco Limited, UK) using a gastight syringe. For the measurement of potential CH_4_ production rates, anoxic conditions were established in the same samples by submerging the soils with degassed H_2_O_deion_ for five days before being transferred into the microcosms. After transferring the soil into the microcosms, they were alternately evacuated to -700 mbar and flushed with N_2_ gas five times to establish anoxic conditions. Three gas samples (12 ml) were taken from the headspace every 24 hours and the atmospheric pressure in the microcosms was re-established by adding 12 ml of N_2_ after each sampling event. CH_4_ and CO_2_ concentrations were measured with a gas chromatograph equipped with a flame ionization detector (FID) and a methanizer to measure CO_2_ (Agilent 7890; Agilent Technologies Inc.). The slope of a linear regression against time was used to calculate gas formation or consumption rates. Three standard gases with known concentrations were used for calculations of the gas concentrations in the gas samples.

### 2.4 DNA extraction and qPCR

DNA was extracted from 0.35 g of soil with the Qiagen DNeasy PowerSoil Kit and stored at −20°C. DNA concentrations were measured on a NanoDrop™ 8000 (Thermo Fischer Scientific). Three groups of methanotrophic bacteria were quantified with three different quantitative PCR assays with a qTower3G (Analytik Jena). A general *pmoA* assay was used to detect a broad spectrum of canonical low affinity methanotrophs (Costello and Lidstrom, 1999), the FOREST assay (Kolb et al., 2003) to quantify USCα specific *pmoA*, and the GAM assay to quantify a USCγ specific *pmoA* (Kolb et al., 2005). Moreover, methanogens were quantified with an *mcrA* specific assay (Angel et al., 2011). To correct for sample specific inhibition in the PCR reaction, the Inhib-Corr Assay was used to calculate an individual inhibition factor for each DNA extract (Degelmann et al., 2010). The qPCR reactions (15 µl) were performed in 96-well plates with innuMIX qPCR Sygreen Sensitive (IST Innuscreen GmbH, Berlin) using a three-step thermal profile with denaturation at 95 °C for 25 s, annealing at assay specific temperature (Table S1) for 20 s, and elongation at 72 °C for 45 s. Bovine serum albumin was added to the master mix to reach a final concentration of 2 ng/µl. Data from qPCR and gas flux measurements is stored in BExIS (http://doi.org/10.17616/R32P9Q; accession number 32025; Volles et al., 2025b).

### 2.5 Statistics

We constructed linear mixed effects models with the lme4 package (Bates et al., 2015) in R (R Core Team, 2024) to test for an effect of the experimental reduction of LUI on the measured variables. To increase the statistical signal, we constructed separate models for each of the soil depths. We constructed an overall model that included all 45 experimental sites, each consisting of a reduced LUI plot and an intensively managed control plot from all three regions. To account for the hierarchical regional and site-level variation, both region and site were modeled as random effects with random intercepts and site identity was nested within region. To identify region-specific effects, each of the three regions was tested with a separate model with only site identity as a random factor with a random intercept. A second model structure was used to test for the difference between the experimentally reduced LUI plots and sites with historically low LUI. Here, only region was a random factor with a random intercept since site level identity did not match the experimental sites. To compare reduced LUI plots with historically low LUI sites on a regional level, a linear model without random effects was used. Model assumptions were visually assessed using residual plots and Q-Q plots. Copy numbers from qPCR were log-transformed to meet the normality assumption for model residuals.

## 3. Results

### 3.1 Methanotrophy and potential CH_4_ oxidation rates (PMORs)

All top- and subsoil samples exhibited uptake of atmospheric CH_4_ with PMORs ranging between 0.08 and 0.97 ng CH_4_ g^−1^ DW hr^−1^ (Figure 1a). In subsoil, PMOR was generally lower than in topsoil (-16.24%; p < .001; Figure S2a). The reduction of LUI had no effect on PMOR in top- and subsoil, neither including all, nor in any of the three individual regions separately. Comparison of reduced LUI plots to historically low LUI sites revealed significant differences with higher PMOR in the topsoil of the historically low LUI sites overall (+ 47.6%; p < .001) and in all three regions (ALB: + 41.8%; p < .001, HAI: + 35.3%; p < .01, SCH: + 58.5%; p < .001, Figure 1a). In subsoil, the effect was significant overall (+ 30.2%; p < .01) and in the region Schorfheide-Chorin (SCH: + 48.6%; p < .001) while it was not in the regions Schwäbische Alb (ALB) and Hainich-Dün (HAI) (Figure S2a). In topsoil samples, PMOR was overall negatively correlated with CO_2_ production while there was an overall positive correlation in subsoils (Figure S5).

**Figure 1:**
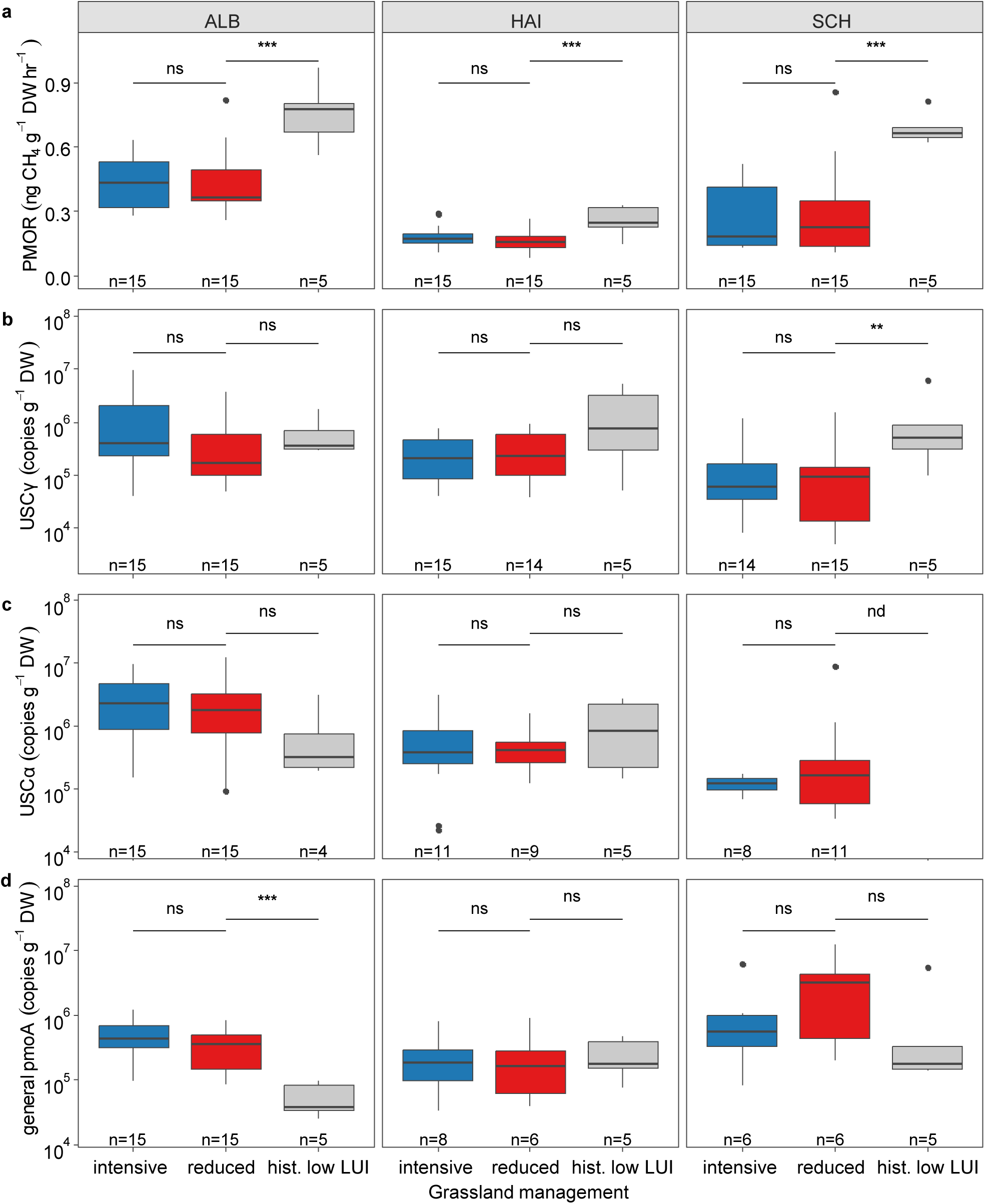
(**a**) Potential methane oxidation rates (PMORs), (**b**) USCγ methanotroph gene abundance, (**c**) USCα methanotroph gene abundance, and (**d**) general *pmoA* gene abundance representing canonical methanotrophs in topsoil (0-10 cm) from different experimental regions of Schwäbische Alb (ALB), Hainich-Dün (HAI) and Schorfheide-Chorin (SCH). Significance codes: p < .05 (*), p < .01 (**), p < .001 (***).

USCγ methanotrophs were detected in 97.6% of all soil samples with gene abundance that ranged from 1.1 × 10^4^ to 9.7 × 10^6^ USCγ-type *pmoA* genes g^-1^ DW (Figure 1b). In the subsoil, USCγ gene abundance was 62% lower than in the topsoil (p < .001; Figure S2b). The experimental reduction of LUI did not significantly affect the overall USCγ gene abundance in both top- and subsoil. Similar to PMOR, there was significantly higher USCγ gene abundance in historically low LUI grassland sites than in experimentally reduced LUI plots (+75%; p< .01; Figure 1b). This increase was significant in the region SCH (+90.7%; p = .01) but not in ALB (+46.5%) and HAI (+68.2%).

USCα methanotrophs were detected in 74.3% of topsoil samples and 60% of subsoil samples with gene abundance ranging from 4.2 × 10^4^ to 1.3 × 10^7^ USCα-type *pmoA* genes g^-1^ DW (Figure 1c). In subsoil (Figure S2c), gene abundance was significantly lower (-78.9%; p < .001) than in topsoil. The experimental LUI reduction had no effect on USCα gene abundance in both soil depths overall and separately in the regions. In contrast to USCγ, USCα gene abundance was similar between experimentally reduced and historically low LUI sites. In the region SCH, historically low LUI sites did not harbor any USCα genes at all (Figure 1c).

Canonical methanotrophs, represented by qPCR targeting general *pmoA* genes were detected in 68% of all samples and followed a region specific distribution (Figure 1d). Gene abundance ranged from 1.8 × 10^5^ to 1.3 × 10^8^ g^-1^ DW and was generally lower in subsoil (-71.1%; p < .001; Figure S2d). The experimental reduction of LUI did not affect the abundance of general *pmoA* representing canonical methanotrophs. In comparison to the experimentally reduced LUI plots, the historically low LUI sites inhabited lower abundance of general *pmoA* overall (-218.8%; p < .01) which was most pronounced in the region ALB (-506.6%; p < .001) (Figure 1d).

### 3.2 CH_4_ Production potential and abundance of methanogens

CH_4_ production was detected in all soil samples with rates that ranged from 0.01 to 0.75 ng CH_4_ g^−1^ DW hr^−1^ (Figure 2a). In subsoil, similar CH_4_ production rates were observed compared to topsoil (Figure S3a). In both top- and subsoil, experimental reduction of LUI had no effect on CH_4_ production potential neither overall nor in any of the regions separately. Comparison between reduced LUI plots and historically low LUI sites showed no difference overall, but an increase on historically low LUI sites in the topsoil in the ALB region.

**Figure 2:**
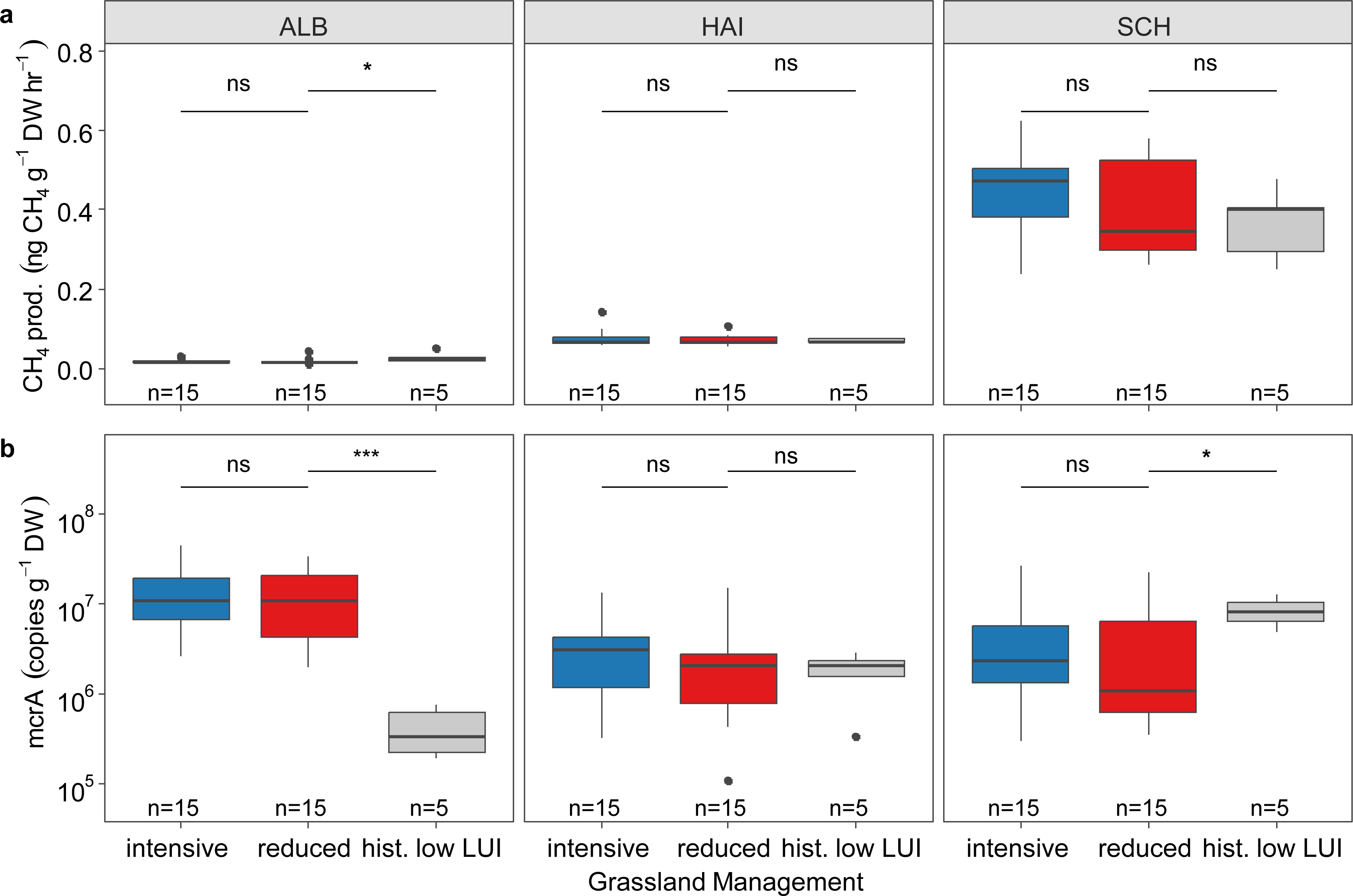
(**a**) Potential methane production rate and (**b**) gene abundance of *mcrA* in topsoil (0-10 cm) of different experimental regions, i.e. Schwäbische Alb (ALB), Hainich-Dün (HAI) and Schorfheide-Chorin (SCH). Significance codes: p < .05 (*), p < .01 (**), p < .001 (***).

The abundance of methanogens (*mcrA* gene copies g^-1^ DW) was higher in topsoil than in subsoil (+ 74.5%; p < .001). The reduction of LUI resulted in a decrease of *mcrA* gene abundance in topsoil overall (-31.4%; p < .05) but not in subsoil (Figure S3b). The comparison to historically low LUI sites revealed no overall effect but a region-specific differentiation (Figure 2b). Historically low LUI sites had lower (-2454.5%, p< .001) *mcrA* gene abundance in the ALB region, a similar one in HAI, and higher abundance in SCH (+ 77.6%; p < .05) as compared to the reduced LUI sites. A similar pattern was observed in the subsoil (Figure S3b).

### 3.3 Relationships between activity and abundance of methanotrophs and methanogens

PMOR and USCγ gene abundance was overall positively correlated. This effect was significant in samples of the regions SCH and ALB, but not in those of HAI (Figure 3a). PMOR and USCα gene abundance was positively correlated overall and in the SCH region but not in HAI and ALB (Figure 3b). Canonical methanotrophs gene abundances represented by the general *pmoA* qPCR assay was not correlated with PMOR in the three regions individually, only taken together an overall effect was observed (Figure 3c). However, general *pmoA* gene abundance was positively correlated with the abundance of *mcrA* and in all regions individually (Figure 4). Moreover, there was a positive correlation between *mcrA* gene abundance and NO_3_ ^−^ concentrations (overall: r = .43; p < .001, ALB: r = .42; p < .001, HAI: r = .46; p < .001, SCH: r = .58; p < .001) and NH ^+^ concentrations (overall: r = .53; p < .001, ALB: r = .38; p < .01, HAI: r = .51; p < .001, SCH: r = .53; p < .001; Figure S6a and b).

**Figure 3:**
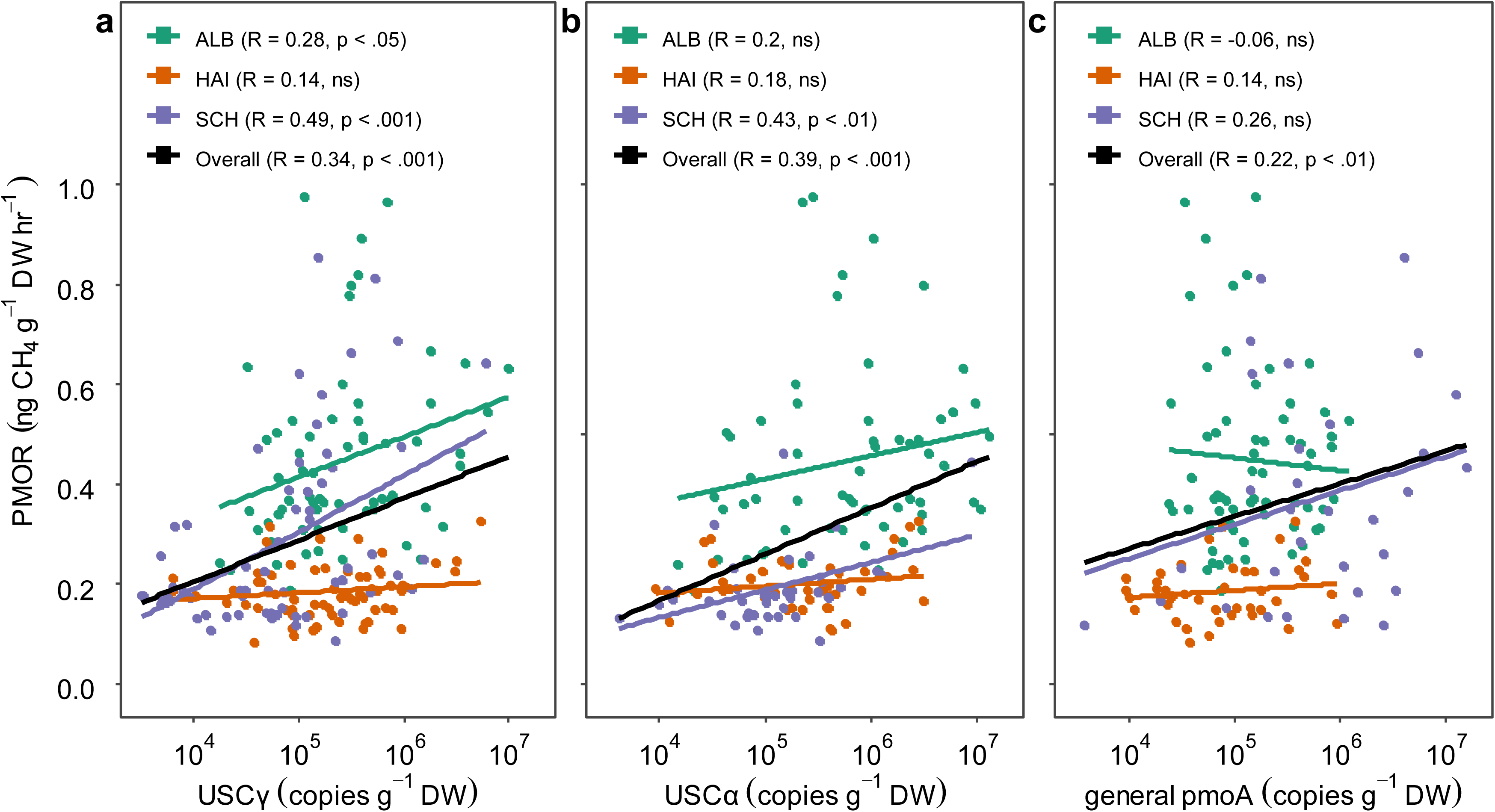
Pearson correlation between potential methane oxidation rate (PMOR) and gene abundance of (**a**) USCγ, (**b**) USCα, (**c**) canonical methanotrophs (general *pmoA*) in top- and subsoil from different experimental regions of Schwäbische Alb (ALB), Hainich (HAI) and Schorfheide-Chorin (SCH).

**Figure 4:**
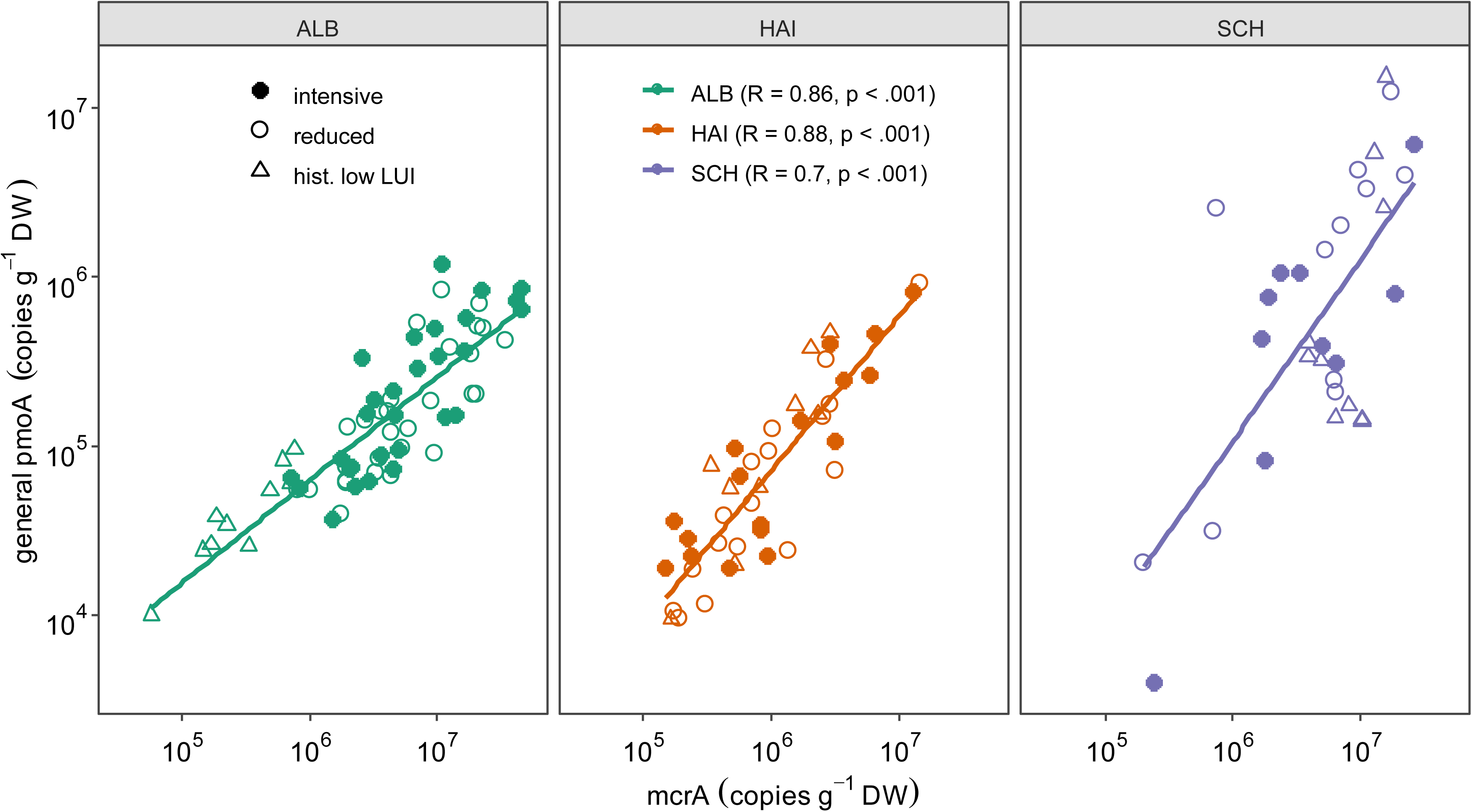
Pearson correlation between gene abundance of canonical methanotrophs (general *pmoA*) and *mcrA* gene abundance in top- and subsoil in the regions of Schwäbische Alb (ALB), Hainich-Dün (HAI), and Schorfheide-Chorin: (SCH).

### 3.4 Soil properties under reduced land-use intensity

Three years of experimental reduction of LUI resulted in decreased soil bulk density (BD) in the regions SCH (-0.17 g/cm^3^; p < .001) and HAI (-0.07 g/cm^3^; p < .05; Figure 5a). Compared to experimentally reduced LUI plots, sites with historically low LUI exhibited an even lower bulk density in the regions SCH (-0.3.7 g cm^-3^; p < .001) and HAI (-0.27 g cm^-3^; p < .001). Bulk density measurements from ALB were inadequate and thus excluded. After three years of experimental LUI reduction, soil water content (SWC) in topsoil was significantly increased (+7.1%; p < .05; Figure S6) and compared to reduced LUI plots, those with historically low LUI had an even higher overall SWC (+ 33.4%; p < .01). SWC in both top- and subsoil was positively correlated with PMOR overall (r = .42; p < .001; Figure S7).

**Figure 5:**
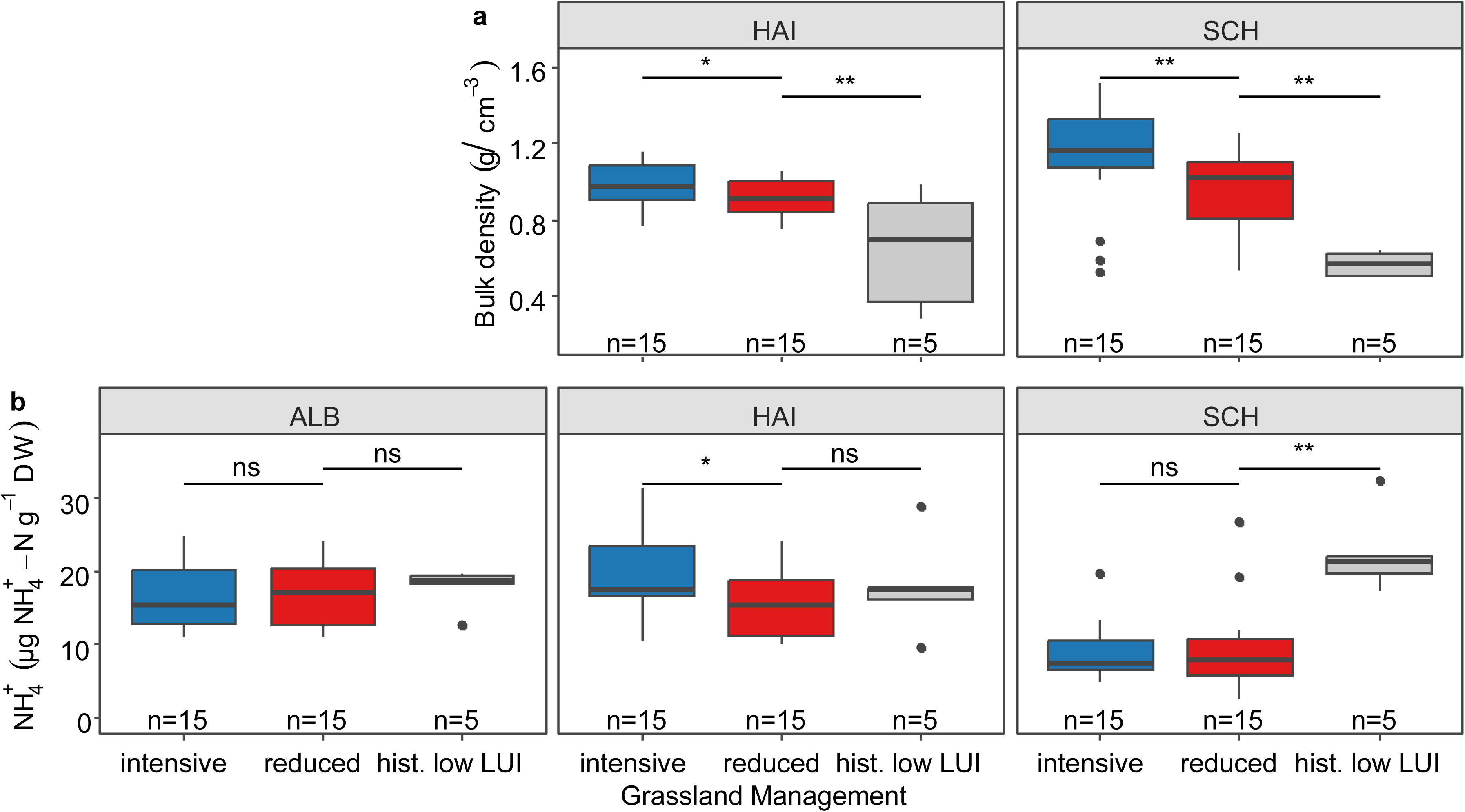
(**a**) Soil bulk density (BD) and (**b**) ammonium concentration in topsoil (0-10 cm) in the regions Schwäbische Alb (ALB), Hainich-Dün (HAI) and Schorfheide-Chorin (SCH). Significance codes: p < .05 (*), p < .01 (**), p < .001 (***).

The experimental reduction of LUI had no effect on other soil physico-chemical parameters in samples from top- and subsoil (pH, N_min_, NO ^−^, C_mic_, EOC, ETN; Figure S8-S13). However, all topsoil samples from historically low LUI sites showed higher values compared to reduced LUI plots: pH +0.6 (p = .01), N_min_ +30.8% (p < .01), NO_3_ ^−^ +37.5% (p < .05), C_mic_ +34.3% (p < .01), EOC +55.5% (p < .01), and ETN +36.1% (p < .01). N_min_ was mainly composed of NH_4_ ^+^. In HAI, NH_4_ ^+^ contents were decreased in the topsoil after three years of experimental LUI reduction (-21%; p <.05). Higher NH_4_ ^+^ values in historically low LUI sites than in experimentally reduced LUI soils (+27.9%, p < .01) were detected in region SCH but not in ALB and HAI (Figure 5b).

## 4. Discussion

### 4.1 Drivers of CH_4_ uptake

CH_4_ uptake into soil is driven by several factors that depend on land-use intensity of grasslands. Bulk density, soil water content and the nitrogen status of the soil are major determinants for the abundance and activity of atmospheric CH_4_-consuming methanotrophs and regulate the response to restoration measures (Wu et al., 2020).

After three years of reduced LUI, soil bulk densities decreased, approaching the even lower level found in historically low LUI grasslands. This is likely due to less soil compaction from heavy machinery and livestock trampling and has been repeatedly documented within the first years of grazing exclosure in grasslands (Wei et al., 2012; Tang et al., 2019) and under secondary succession (McDaniel et al., 2019). Bulk density has been identified as a major driver of atmospheric CH_4_ oxidation in grasslands with legacy effects even after sieving and re-compacting in incubations (Täumer et al., 2021; Sitaula et al., 2000). Higher soil porosity and less soil compaction may increase gas diffusion rate and thus, increase substrate (CH_4_) and terminal electron acceptor (O_2_) availability for atmospheric CH_4_ consuming methanotrophs. However, in 24 days of increased substrate availability, no changes in the methanotroph communities were observed in soil incubations (Malghani et al., 2016). A possible mechanism of CH_4_ sink recovery following a bulk density decrease might however be a long-term process as facultative atmospheric methanotrophs may use other carbon sources (Tveit et al., 2019) and have slow growth rates. Besides bulk density, the soil water content (SWC) is another key factor controlling diffusion rate and CH_4_ oxidation in soils (Täumer et al., 2022). To normalize for this highly dynamic factor over time, we equalized SWC between the intensive and reduced LUI treatment. However, in the field, reduced LUI plots had higher SWC than intensive ones, approaching almost SWC contents of historically low LUI site levels. Higher SWC in reduced and historically low LUI grasslands might be related to changes in evaporation. Grazing of commonly decreases water content of grassland soils due to reduced canopy cover and increased infiltration by livestock trampling (Niu et al., 2025; Gifford and Hawkins, 1978). This might have led to changes of *in situ* fluxes and methanotrophic activity that are not covered in our incubations for PMOR measurements or by DNA-based methanotroph abundance measurements. Although fertilization and grazing were omitted, we observed no significant changes in soil N_min_ concentrations, except for an NH_4_ ^+^-driven reduction on reduced LUI plots in HAI. Notably, this did not lead to changes in CH_4_ oxidation, so that a short-term recovery from possible inhibitory effects of NH_4_ ^+^ by competition for active sites of CH_4_ monooxygenase is unlikely (Schnell and King, 1994). Both, increased and reoccurring application of NH_4_ ^+^ are described to critically reduce CH_4_ oxidation in soils while weakening the community structure, selecting for NH_4_ ^+^-resistant methanotrophs (Ho et al., 2020; López et al., 2019; Lim et al., 2024), and thus substantially delaying recovery (Lim et al., 2024). Such shifts might be partly reflected in lower abundance of USCγ in the topsoil of reduced LUI plots compared to historically low LUI sites overall and in region SCH.

All other soil parameters that we have studied showed no response to the reduction of LUI. Since, these parameters were different on the historically low LUI sites, we consider the status of the recovery processes as too early and minor to affect the CH_4_ sink function. Moreover, none of the other parameters that we analyzed was consistently different on historically low LUI sites throughout the three regions. This indicates that CH_4_ sink recovery may be driven by other, non-studied single factors or that rather a combination of several environmental factors is necessary.

### 4.2 Methanotrophs mediating grassland atmospheric CH_4_ sink recovery

USCγ was detected in 97.6% of top- and subsoil samples which is in line with a previous study in the same regions (Täumer et al., 2021) and for grasslands on a global scale (Deng et al., 2019; Kolb, 2009). Moreover, USCγ dominates the active methanotroph community in *pmoA* transcripts from soil incubations of temperate grasslands under atmospheric conditions (Liu et al., 2022). USCγ dominates in neutral to alkaline soils, where several abundance and activity patterns have been described, but it still misses cultured representatives that would inform about more specific aspects of its ecophysiology and ecological niche (Deng et al., 2019; Knief, 2015). Our study found a 75% higher USCγ gene abundance in historically low LUI sites compared to those with three years of LUI reduction, matching the pattern observed in PMOR measurements. From this finding, we conclude that USCγ methanotrophs play a key role for the recovery of the CH_4_ sink function of grasslands.

The other group of atmospheric CH_4_ consuming methanotrophs, USCα, was less frequently abundant. This is consistent with previous work that found USCα to rather dominate in forest and acidic soils (Kolb, 2009; Knief, 2015; Täumer et al., 2021). USCα is commonly absent in agricultural soils and its abundance and biomass is very sensitive to disturbances by soil management practices such as ploughing (Maxfield et al., 2008; Shrestha et al., 2012). Interestingly, some studies reported re-emergence of USCα after disturbance (Ho et al., 2011; Levine et al., 2011). USCα abundance correlated positively with PMOR overall even in the region SCH where it was not detected in historically low LUI sites at all. Since PMORs were greater on grassland sites with historically low LUI consistently in all regions, our results suggest that USCα plays a minor role in the short-term recovery of the CH_4_ sink of temperate grasslands but may exert additive effects in atmospheric CH_4_ oxidation.

### 4.3 CH_4_ filter function of grassland under reduced land use intensity

The general general *pmoA* assay used in our study, combines a broad spectrum of methanotrophs that oxidize CH_4_ at varying concentrations. However, this qPCR assay largely discriminates against USCα and γ and thus reflects rather the abundance of canonical methanotrophs with low substrate affinity (Kolb et al., 2003; Knief, 2015). This group of methanotrophic microorganisms oxidizes CH_4_ at high concentrations larger than 600 ppmv, which is usually produced by methanogens in anoxic compartments within the soil, thus mediating a filter function for CH_4_ before it reaches the atmosphere (Conrad, 1996; Knief, 2019). We found a remarkably strong positive linear relationship between the abundance of canonical methanotroph *pmoA* genes and methanogen *mcrA* genes in all tested regions. The decline in gene abundance from topsoil to subsoil was consistent across all quantified groups. This is expected for atmospheric CH_4_ consuming and canonical methanotrophs (McDaniel et al., 2021; Wang et al., 2023b) but not for the strictly anaerobic and O_2_ sensitive methanogens (Liu and Whitman, 2008). In general, the total amount of microorganisms is higher in the topsoil than in subsoil (Taylor et al., 2002). So even if the *mcrA* genes occur more frequently in the topsoil relative abundance is higher in subsoil which is supported by a greater *mcrA* to C_mic_ in the subsoils we analyzed. In fact, methanogens have been shown to be ubiquitous in aerated upland soils and can be activated under anoxic conditions (Angel et al., 2012).

The calculated *pmoA* to *mcrA* gene ratios together with the remarkably strong *pmoA* to *mcrA* correlations be informing about the soil CH_4_ filter function. For instance, in both SCH and HAI *mcrA* gene abundances were on average approximately 10 times higher than *pmoA* gene abundances. If gene abundance reflects organism abundance and our soils ecosystems have reached the carrying capacity for canonical methanotrophs, one might speculate that an efficient CH_4_ filter requires approximately one methanotroph cell per ten methanogens. However, this assumption does not take into account bioenergetics and soil edaphic factors as well as the potentially varying distribution of methanotrophs and methanogens within the soils matrix. *pmoA*/*mcrA* gene ratios could nevertheless prove useful for assessing the CH_4_ sink or source status of grasslands (Täumer et al., 2022).

The activity and abundance of the soil bound CH_4_ filter community are multi-faceted influenced. Besides soil type, soil moisture, bulk density, vegetation and air temperature and precipitation are key factors for the composition of the soil microbiomes (Hansen et al., 2024; Wang et al., 2023a). Long-term manure application with input of methanogens originating from cattle rumen can shift the methanotroph to methanogen ratio and thus, reduce the CH_4_ filter function until arable soils turn from sink to source (Gattinger et al., 2007). In a meta-analysis, nitrogen application generally reduced *mcrA* abundance in grasslands (Liu et al., 2024) which is not supported by our study, which found a positive correlation between *mcrA* and both NH_4_ ^+^ and NO ^−-^concentration. Interestingly, our results revealed a decrease of *mcrA* abundance through LUI reduction including the abandonment of grazing and fertilization. This might be interpreted as a first indicator for a beginning long-term restoration of the CH_4_ sink potential.

However, regional differences in *mcrA* and canonical metanotrophs *pmoA* abundance on historically low LUI sites point towards a divergent sensitivity of the CH_4_ filter function to LUI in the studied regions. In the ALB region, historically low LUI sites had lower methanogen abundances, which could result in lower CH_4_ production and, consequently, less substrate for the reduced number of canonical methanotrophs observed. The low CH_4_ production potential may thus be critical for the abundances of canonical methanotrophs in the ALB region. While the CH_4_ filter function of grasslands in the ALB region appears to be sensitive to LUI, it is less sensitive in organic rich soils with higher CH_4_ production potential of HAI and especially SCH. Hence, the introduction of methanogens and their organic substrates through manure application may only lead to an increased abundance of canonical methanotrophs in soils with naturally low levels of methanogenesis.

### 4.4 The time dependency of the CH_4_ sink recovery in grasslands

Omitting fertilization and grazing, as well as minimizing mowing frequency for three years did not affect atmospheric potential CH_4_ oxidation rates (PMORs) of the investigated grassland soils. Sites with historically low LUI had 47.6% higher PMORs compared to the experimentally reduced LUI grassland sites, representing a level that may be achieved through long-term reduction of grassland management practices. This corresponds to previous work within the same regions confirming 40% higher PMORs on low than on high LUI sites (Täumer et al., 2021). While the negative effects of heavy grazing on the CH_4_ sink function are well described (Tang et al., 2019), most existing studies on the recovery from intensive management have investigated grazing exclosure only on single sites, resulting in fragmented and inconsistent finding. For example, it has been shown that four years of grazing exclosure had, depending on soil type, no or even negative effects on the CH_4_ sink function in a prairie grassland (Thomas et al., 2017). In contrast, in an alpine grassland of the Tibetan plateau, a 34% increase of CH_4_ uptake already after 4 years of exclosure while another study found no effect even after 6 years (Wei et al., 2012; Wang et al., 2023c). In Inner Mongolia, 33 years of grazing exclosure resulted in 42% higher CH_4_-uptake (Pan et al., 2021) corresponding well to the 47.6% PMOR increase on historically low LUI sites in our study. (Liu et al., 2022) investigated the effect of a four-year grazing exclosure at 11 different grassland sites spanning 800 km and observed a decreasing effect on the CH_4_ sink function in desert grasslands and no effect in meadow grasslands.

Among these studies about grazing exclosure in grasslands, other studies investigated the CH_4_ dynamics along secondary successional gradients. No *in situ* effect of pasture abandonment was found after 20 years (Keller and Reiners, 1994) and 80 years for complete CH_4_ sink recovery were extrapolated by Levine et al. (2011) being supported by different meta-analyses (McDaniel et al., 2019; Wu et al., 2020). Disturbances of methanotrophs can be differentiated into sporadic and recurring events that take days to weeks to recover from (Lim et al., 2024). However, prolonged or compounded disturbances take years to decades to recover by altering the methanotroph biodiversity. Our results support that intensive grazing as well as mowing together with fertilization have caused such a compounded and prolonged disturbance of the CH_4_ sink function in the grasslands that we have studied.

It has been shown that in heavily degraded grasslands, shallow ploughing but not harrowing can increase the CH_4_ sink function more than natural recovery (Liu et al., 2021). This effect was attributed to increased belowground plant biomass, litter accumulation and soil properties after mechanical soil disturbances. Also plant species identity and diversity have been shown to influence CH_4_-oxidation in grassland ecosystems (Praeg et al., 2017; Ström et al., 2005). However, changes of the plant community are a long-term process as well due to delayed immigration events of new species or delayed extinction events following restricted nutrient availability (Jackson and Sax, 2010; Andraczek et al., 2023). Therefore, further research is required on how to improve the rate of recovery of the CH_4_ sink to increase the mitigation function of grasslands for atmospheric CH_4_.

## 5. Conclusions

In the context of climate change, where soils are expected to contribute to greenhouse gas mitigation (Paustian et al., 2016), the question arises if CH_4_ sink restoration can be achieved through extensification of grassland management. Our study shows that the CH_4_ sink potential of temperate grasslands does not recover within three years of extensification. This finding is highly replicated and consistent across pedoclimatic regions and soil types. Neither methanotroph activity nor abundance changed following three years without fertilization, grazing, and only one mowing event per year. However, a decrease in *mcrA* abundance might signal early shifts in the CH_4_ filter function. Bulk density and soil water content (SWC) also responded to experimental reduction of LUI, aligning with levels in historically low LUI grasslands and suggesting they could be major environmental determinants mediating CH_4_ sink recovery. Despite this, historically low LUI sites consistently exhibited higher PMORs, suggesting CH_4_ potential might be achieved through long-term extensification. The correlation between USCγ abundance and PMORs indicates its key role in CH_4_ sink function and its recovery, while USCα may contribute additional effects. Overall, the results point to the need to study the effects of reducing land use intensity over time, with the medium term likely to be the important transitional period for elucidating the underlying mechanisms that restore the CH_4_ sink function of grassland soils.

## Acknowledgements

This project was funded by DFG Priority Program 1374, “Infrastructure-Biodiversity-Exploratories”, project number 512405836 (acronym MetGrass). We thank the managers of the three Biodiversity Exploratories Julia Bass, Max Müller, Anna K. Franke, Robert Künast, Melissa Jüds, Franca Marian, and all former managers for their work in maintaining the plot and project infrastructure; the many people involved in the coordinated soil sampling campaign 2023; Victoria Grießmeier, for giving support through the central office, Andreas Ostrowski for managing the central data base and Markus Fischer, Eduard Linsenmair, Dominik Hessenmöller, Daniel Prati, Ingo Schöning, François Buscot, Ernst-Detlef Schulze, Wolfgang W. Weisser and the late Elisabeth Kalko for their role in setting up the Biodiversity Exploratories project. Field work permits were issued by the responsible state environmental offices of Baden-Württemberg, Thüringen, and Brandenburg.

## Data availability

This work is based on data elaborated by a project of the Biodiversity Exploratories program (DFG Priority Program 1374). The datasets are publicly available in the Biodiversity Exploratories Information System (http://doi.org/10.17616/R32P9Q) under the accession numbers 32025 (abundance and activity data), 32036 (soil physico-chemical data), and 26207 (pF data of soils). The datasets are listed in the references section.

## Glossary

CH_4_: Methane
CO_2_: Carbon dioxide
pMMO: Particulate methane monooxygenase
*pmoA*: Gene encoding the alpha subunit of pMMO
*mcrA*: Gene used as a biomarker for methanogens
USCα: Upland soil cluster alpha
USCγ: Upland soil cluster gamma
PMOR: Potential methane oxidation rate
LUI: Land-use intensity index
SCH: Schorfheide-Chorin (study region)
HAI: Hainich-Dün (study region)
ALB: Schwäbische Alb (study region)
REX: Reduced land-use intensity experiment
BExIS: Biodiversity Exploratories information system
SWC: Soil water content
C_mic_: Microbial biomass carbon
EOC: Extractable organic carbon
ETN: Total extractable nitrogen
BD: Bulk density
FID: Flame ionization detector
qPCR: Quantitative polymerase chain reaction

