## Supplementary Material for "Methane sink function of grassland soil microbiomes - negative effects of intensive management persist three years after land-use extensification"

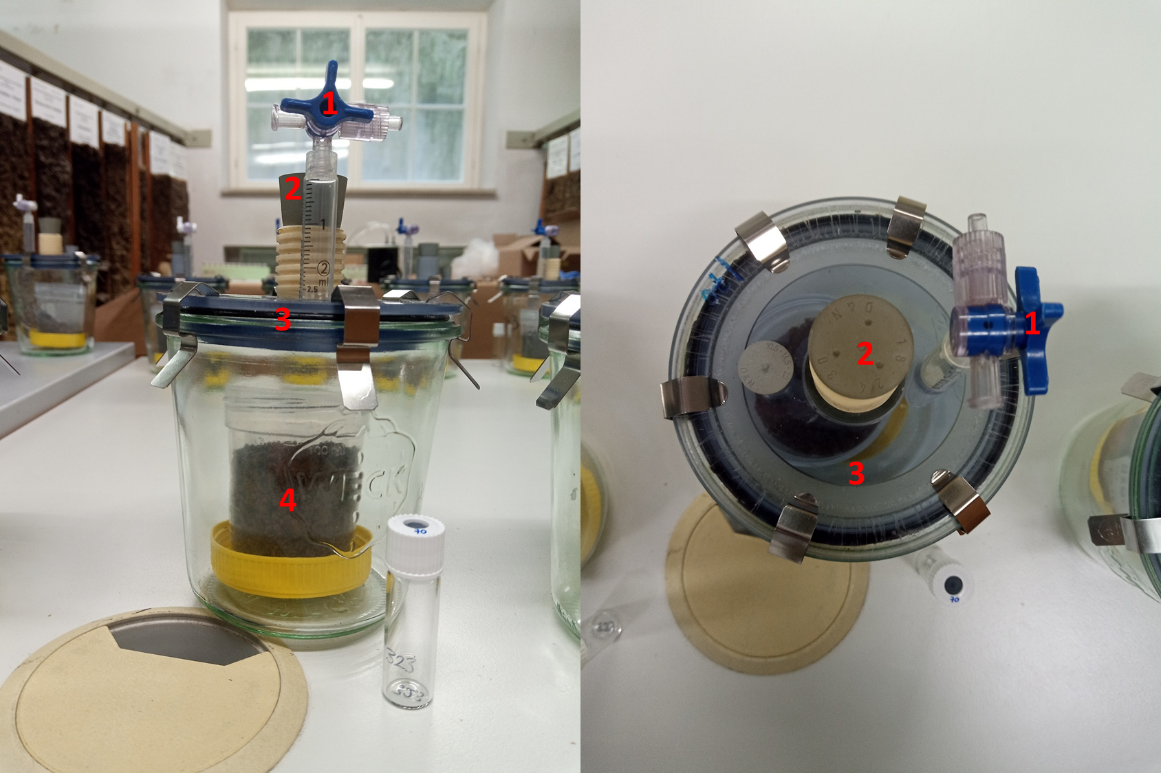

Figure S1: Mesocosm set-up. Gas samples were taken via the valve (1) on the modified gastight lid (3). The aperture (2) allows a simpler gas exchange during ventilation of the microcosms. The soil samples were placed in a plastic beaker (4).

Table S1 qPCR and PCR assays and assay conditions used to quantify functional genes of methanotrophic bacteria methanogenic archaea.

| Assay | Primer | sequence (5’ – 3’) | Amplicon  length (bp) | Primer concentration | | Annealing temperature/time | Data acquisition temperature | Reference |
| --- | --- | --- | --- | --- | --- | --- | --- | --- |
| General pmoA | A189f/ mb661r | GGNGACTGGGACTTCTGG/  CCGGMGCAACGTCYTTACC | 491 | 0.5 µM | 63(25*) + Touchdown (68 10*Δ-0,5) °C/20 s | | 72 °C | (Costello & Lidstrom, 1999) |
| FOREST  (USC-α) | A189f/ forest675r | GGNGACTGGGACTTCTGG /  CCYACSACATCCTTACCGAA | 506 | 0.5 µM | 64 (25*) Touchdown 69 10*Δ-0,5) °C/20 s | | 72 °C | (Kolb, Knief, Stubner, & Conrad, 2003) |
| GAM  (USC–γ) | A189f/ gam634r | GGNGACTGGGACTTCTGG/  ACGAAGCGGATGTACTCGGG | 465 | 0.8 µM | 68 °C/20 s | | 72 °C | (Kolb, Knief, Dunfield, & Conrad, 2005) |
| Inhib CORR | T7f /  M13r | TAATACGACTCACTATAGGG/  CAGGAAACAGCTATGAC | 221 | 0.5 µM | 58 °C/20 s | | 72 °C | (Degelmann, 2010) |
| mcrA | Mlas-mod/  mcrA-rev | GGYGGTGTMGGDTTCACMCARTA/; CGTTCATBGCGTAGTTVGGRTAGT | 470 | 0.5 µM | 55 °C/45 s | | 72 °C | (Angel et al., 2011) |

Costello, A. M., & Lidstrom, M. E. (1999). Molecular characterization of functional and phylogenetic genes from natural populations of methanotrophs in lake sediments. *Appl Environ Microbiol*, *65*(11), 5066–5074.

Kolb, S., Knief, C., Dunfield, P. F., & Conrad, R. (2005). Abundance and activity of uncultured methanotrophic bacteria involved in the consumption of atmospheric methane in two forest soils. *Environmental Microbiology*, *7*(8), 1150–1161. https://doi.org/10.1111/j.1462-2920.2005.00791.x

Kolb, S., Knief, C., Stubner, S., & Conrad, R. (2003). Quantitative Detection of Methanotrophs in Soil by Novel pmoA -Targeted Real-Time PCR Assays. *Applied and Environmental Microbiology*, *69*(5), 2423–2429. https://doi.org/10.1128/AEM.69.5.2423

Degelmann, D. M., Borken, W., Drake, H. L., & Kolb, S. (2010). Different Atmospheric Methane-Oxidizing Communities in European Beech and Norway Spruce Soils. *Applied and Environmental Microbiology, 76(10),* 3228-3235. https:// doi:10.1128/AEM.02730-09

Angel, R., Matthies, D., & Conrad, R. (2011). Activation of methanogenesis in arid biological soil crusts despite the presence of oxygen. PLoS ONE, 6(5). https://doi.org/10.1371/journal.pone.0020

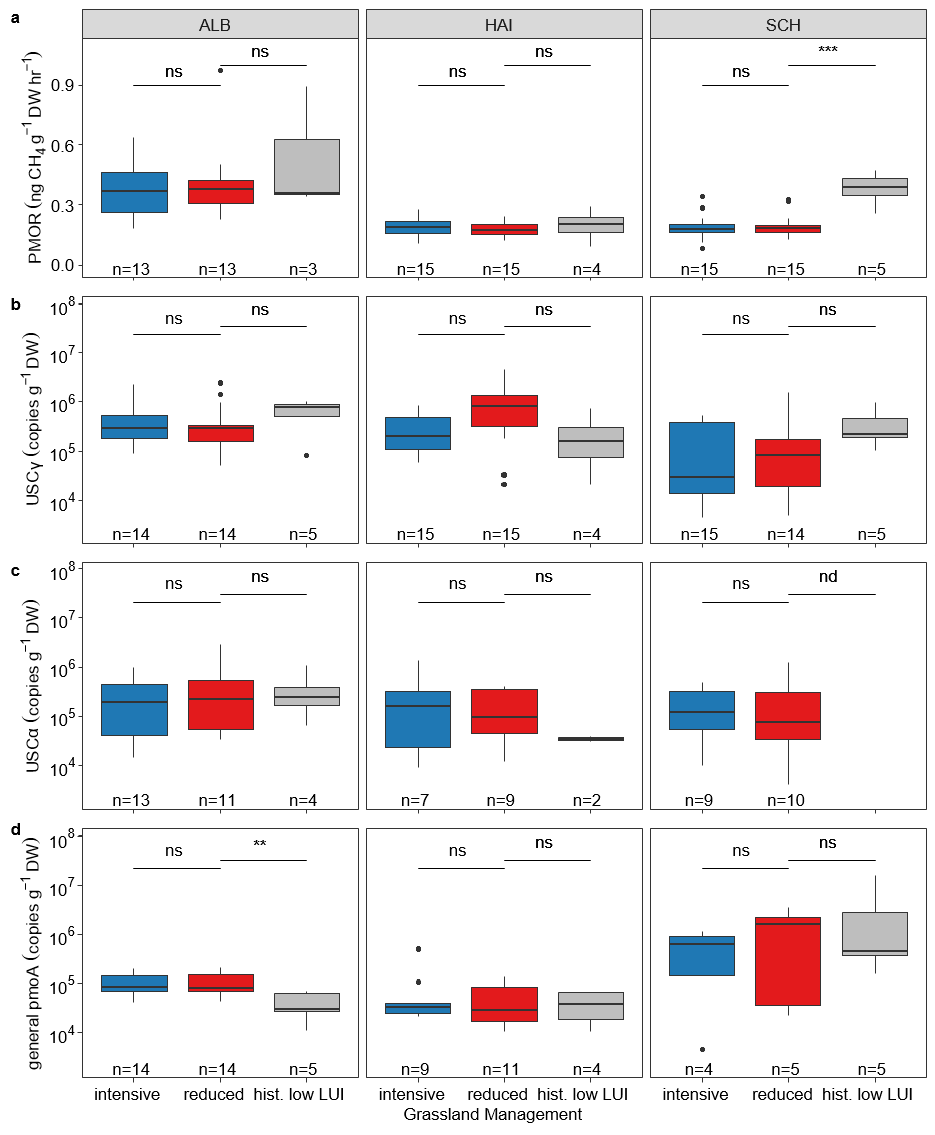

Figure S2: (**a**) Potential methane oxidation rates (PMOR), (**b**) USCγ methanotrophs, (**c**) USCα methanotrophs, (**d**) general pmoA abundance in subsoil and in different experimental regions of Schwäbische Alb (ALB), Hainich (HAI) and Schorfheide (SCH). Significance codes: p < .05 (*), p < .01 (**), p < .01 (***). Intensive vs reduced are compared with a different linear model structure than reduced vs hist. low LUI.

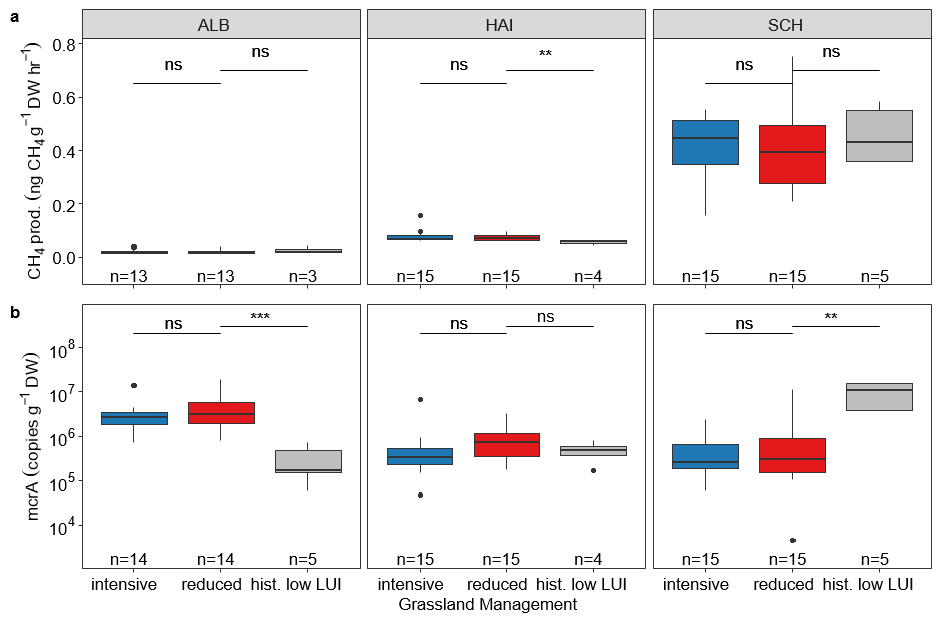

Figure S3: (**a**) Potential methane production and (**b**) abundance of mcrA in subsoil in different experimental regions of Schwäbische Alb (ALB), Hainich (HAI) and Schorfheide (SCH). Significance codes: p < .05 (*), p < .01 (**), p < .01 (***). Intensive vs reduced are compared with a different linear model structure than reduced vs hist. low LUI.

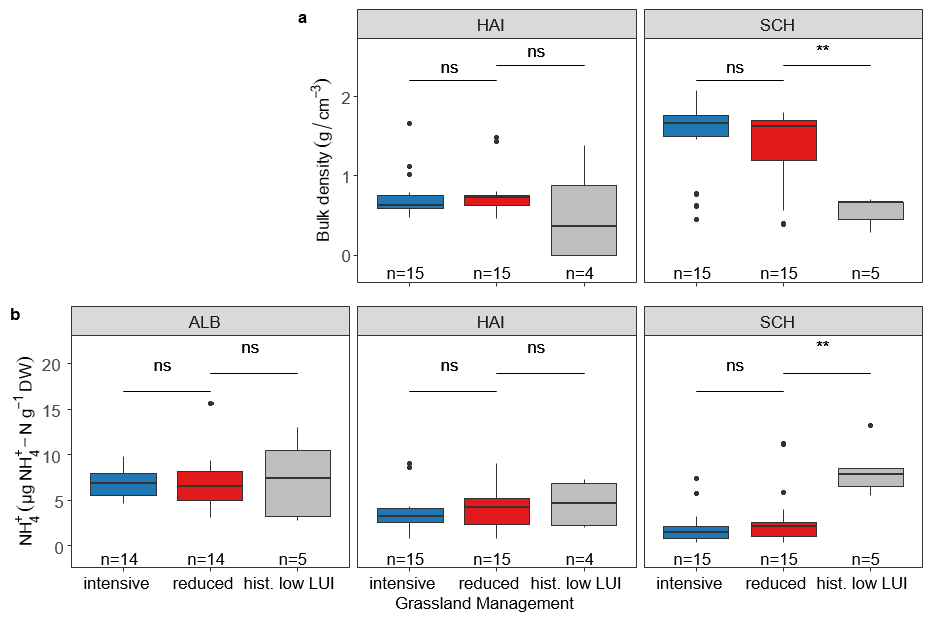

Figure S4: (**a**) Bulk density and Ammonium (**b**) in subsoil (0-10 cm) under de-intensification in the regions of Schwäbische Alb (ALB), Hainich (HAI) and Schorfheide (SCH). Intensive vs reduced are compared with a different linear model structure than reduced vs hist. low LUI.

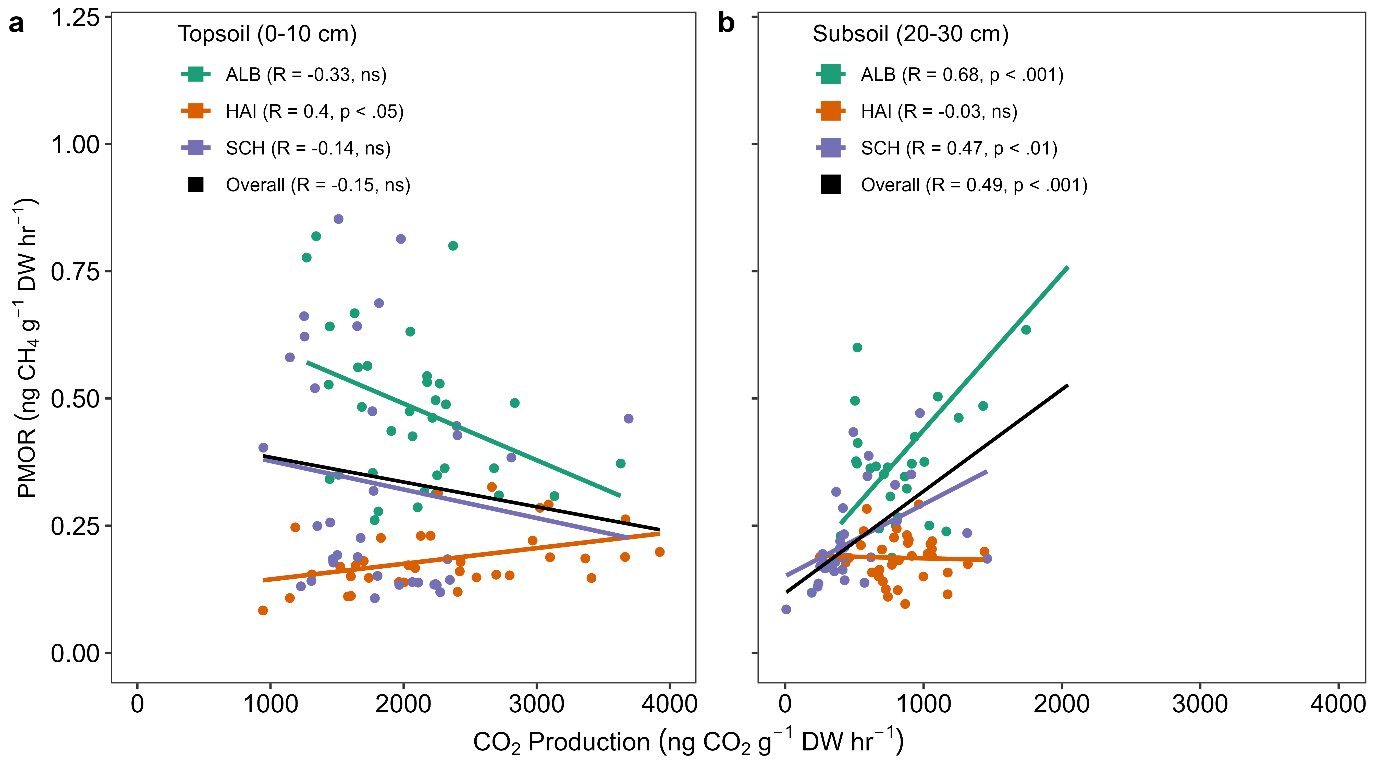

Figure S5: Correlation between potential methane oxidation rate (PMOR) and potential CO_2_ production rate in (**a**) topsoil (0-10 cm) and (**b**) subsoil (20 -30 cm) in microcosm experiment under atmospheric mixing ratios.

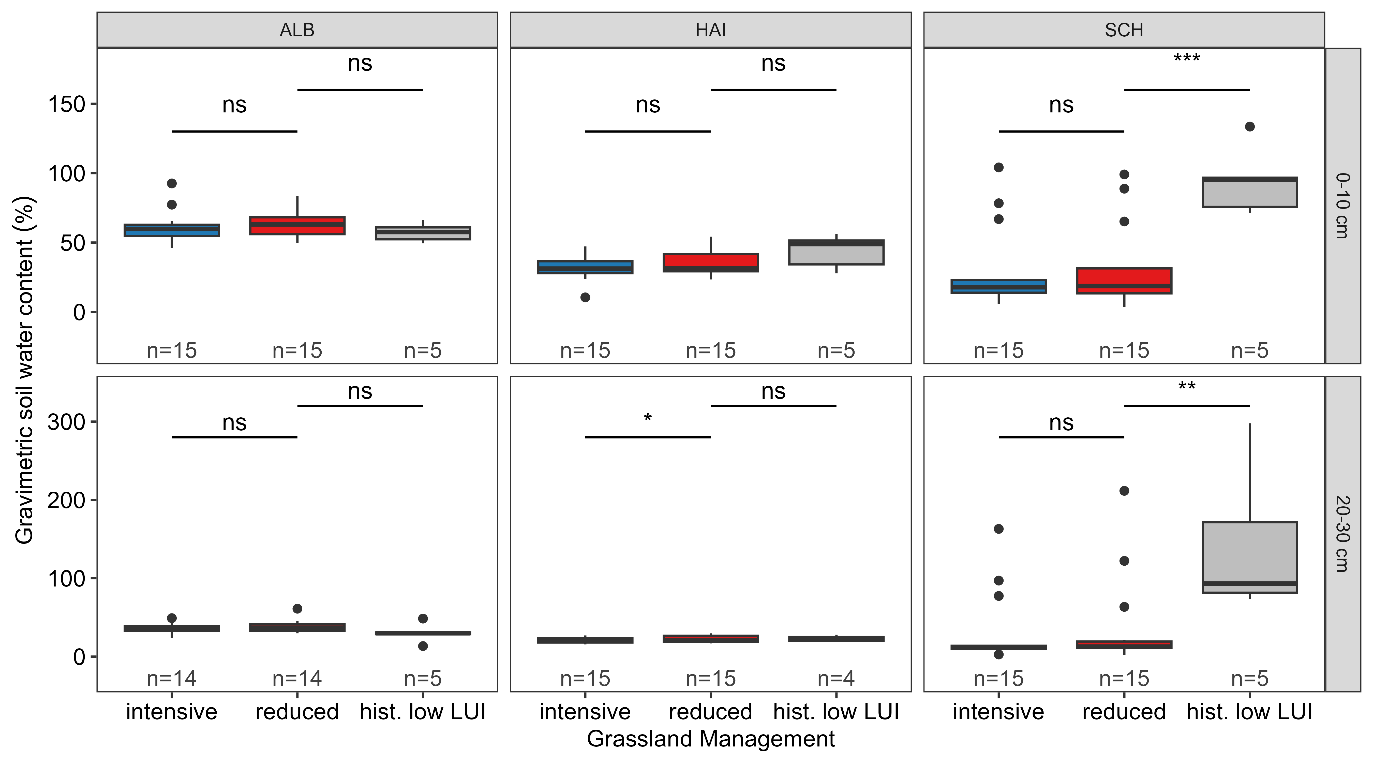

Figure S6: Gravimetric soil water content (SWC) in top- and subsoil in in the regions of Schwäbische Alb (ALB), Hainich (HAI) and Schorfheide (SCH).

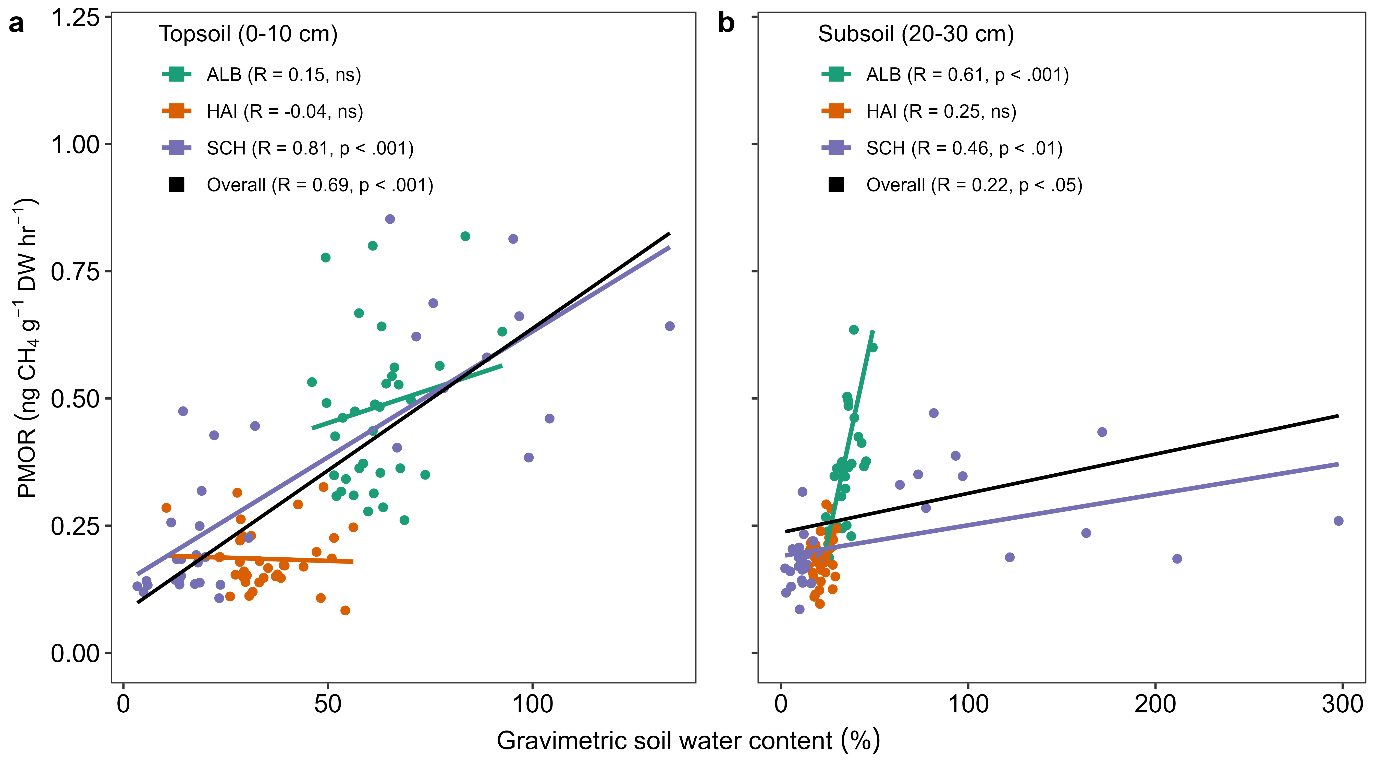

Figure S7: Correlation between potential methane oxidation rate (PMOR) and gravimetric soil water content (SWC) in (**a**) topsoil (0-10 cm) and (**b**) subsoil (20 -30 cm).

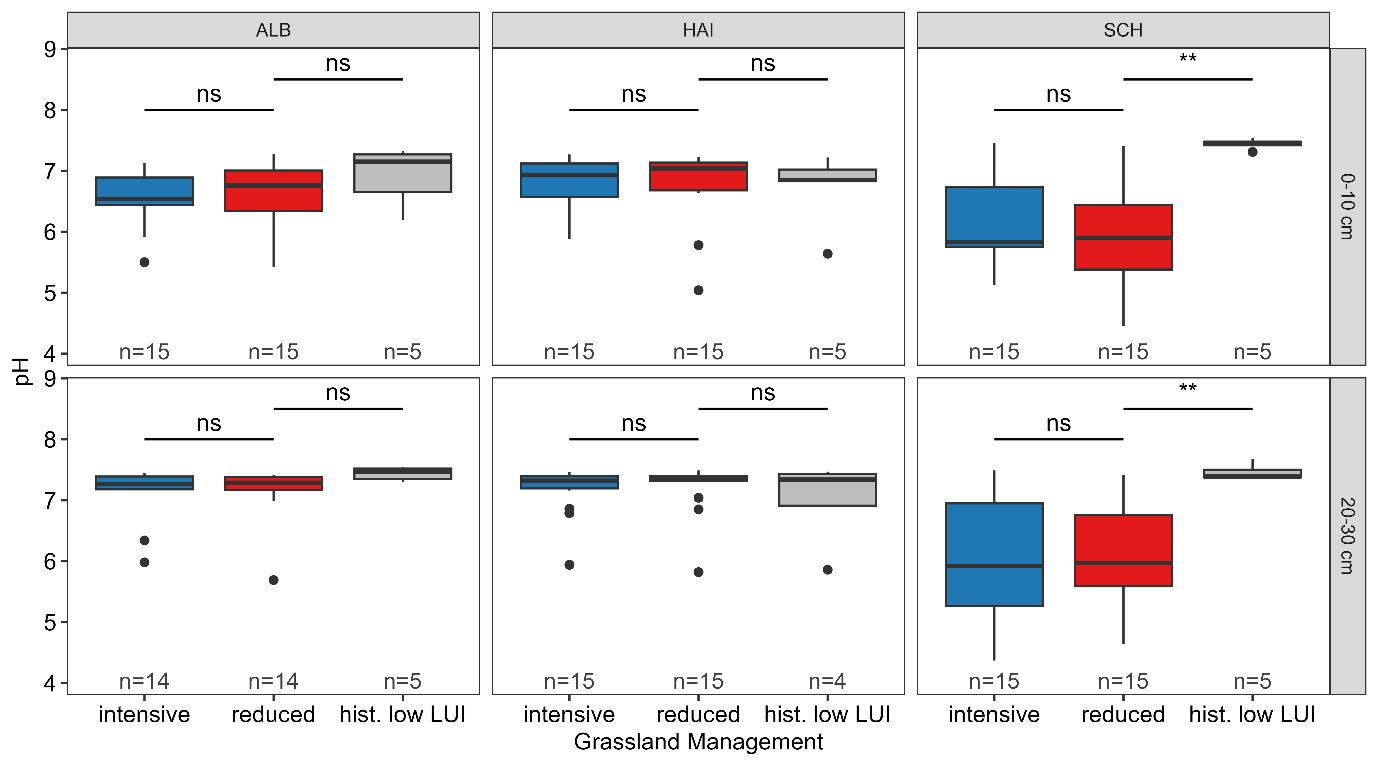

Figure S8: pH in top- and subsoil in in the regions of Schwäbische Alb (ALB), Hainich (HAI) and Schorfheide (SCH).

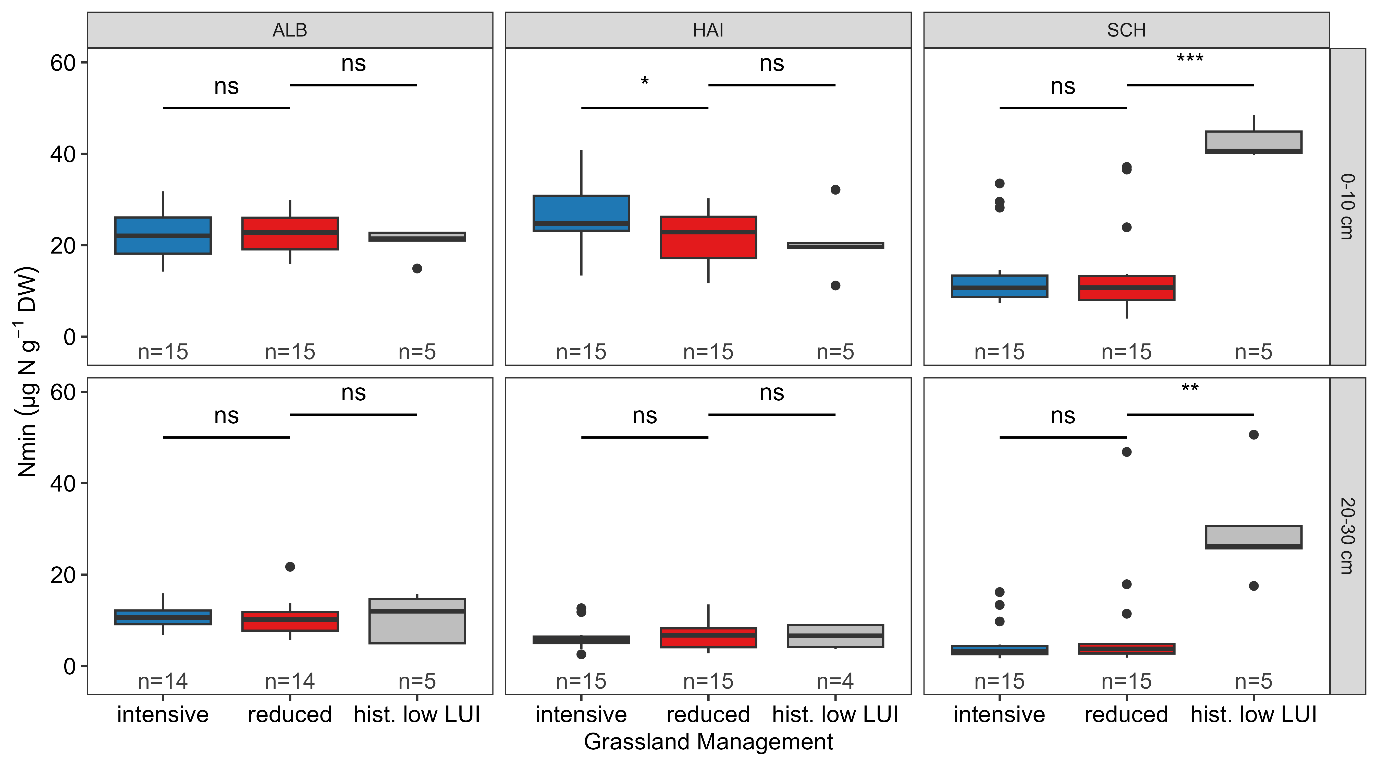

Figure S9: N_min_ in top- and subsoil in in the regions of Schwäbische Alb (ALB), Hainich (HAI) and Schorfheide (SCH).

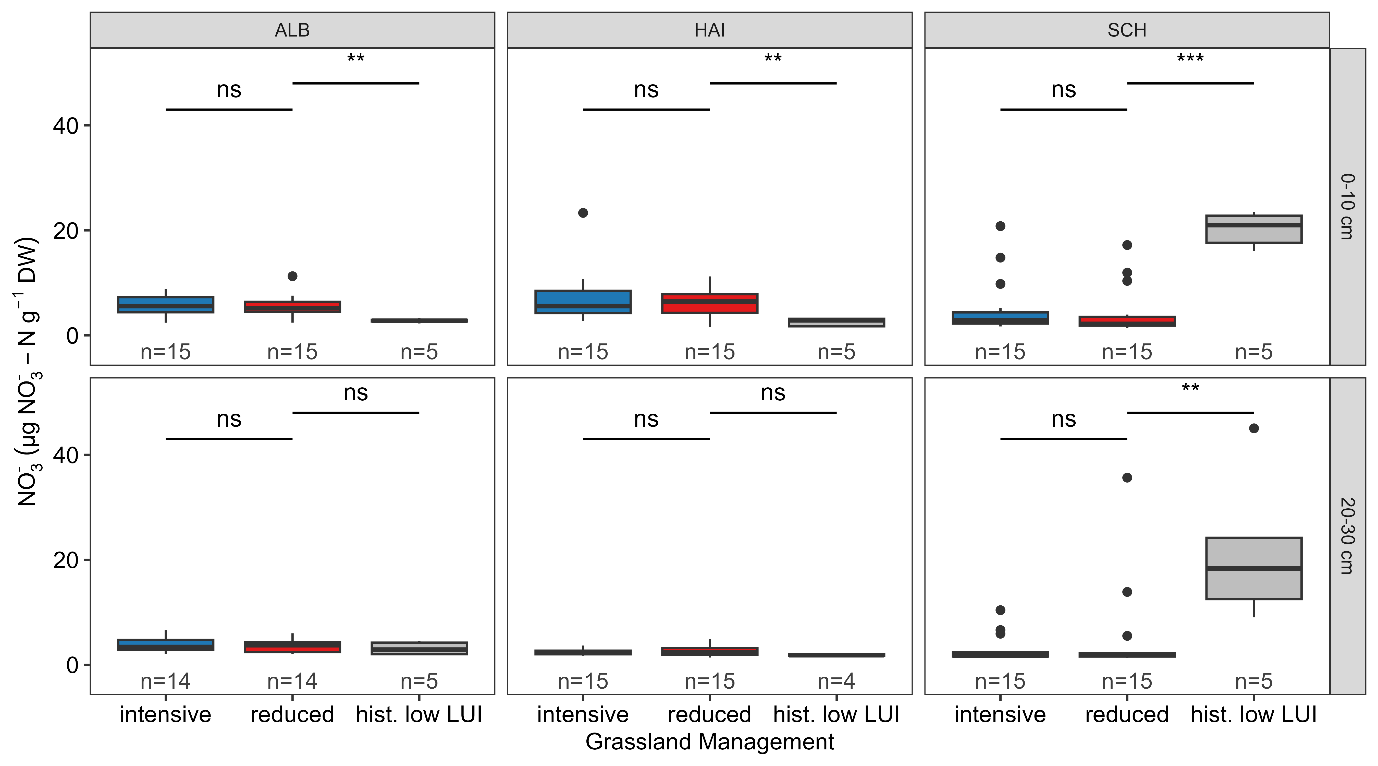

Figure S10: Nitrate in top- and subsoil in in the regions of Schwäbische Alb (ALB), Hainich (HAI) and Schorfheide (SCH).

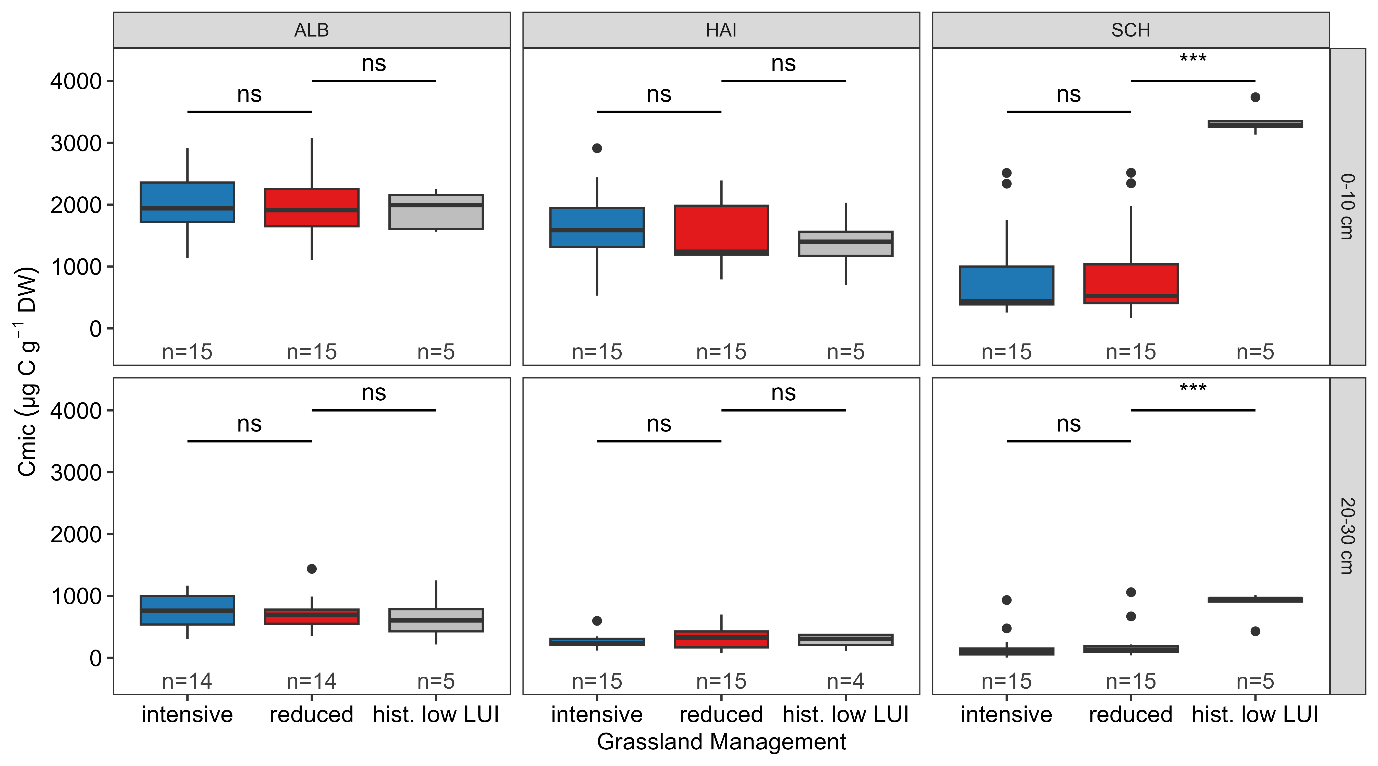

Figure S11: C_mic_ in top- and subsoil in in the regions of Schwäbische Alb (ALB), Hainich (HAI) and Schorfheide (SCH).

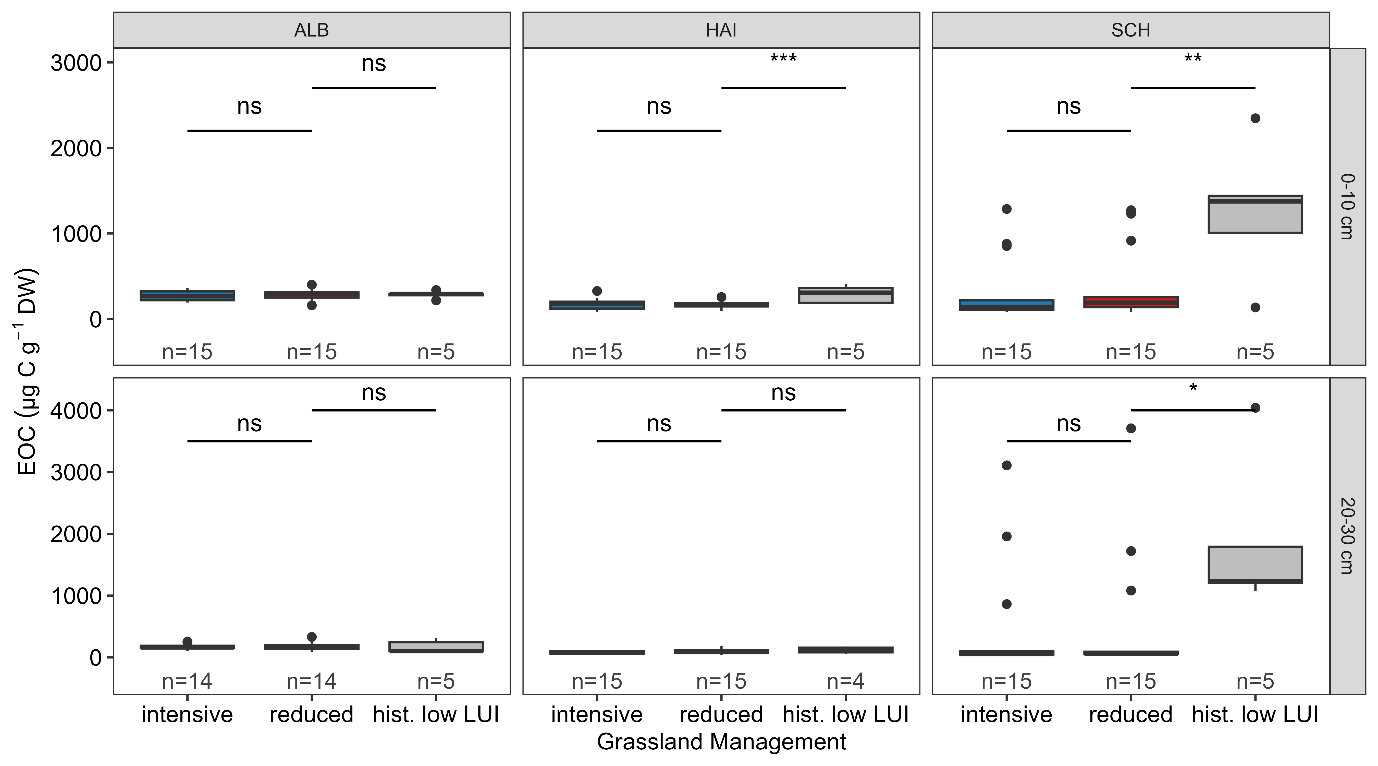

Figure S12: Extractable organic carbon (EOC) in top- and subsoil in in the regions of Schwäbische Alb (ALB), Hainich (HAI) and Schorfheide (SCH).

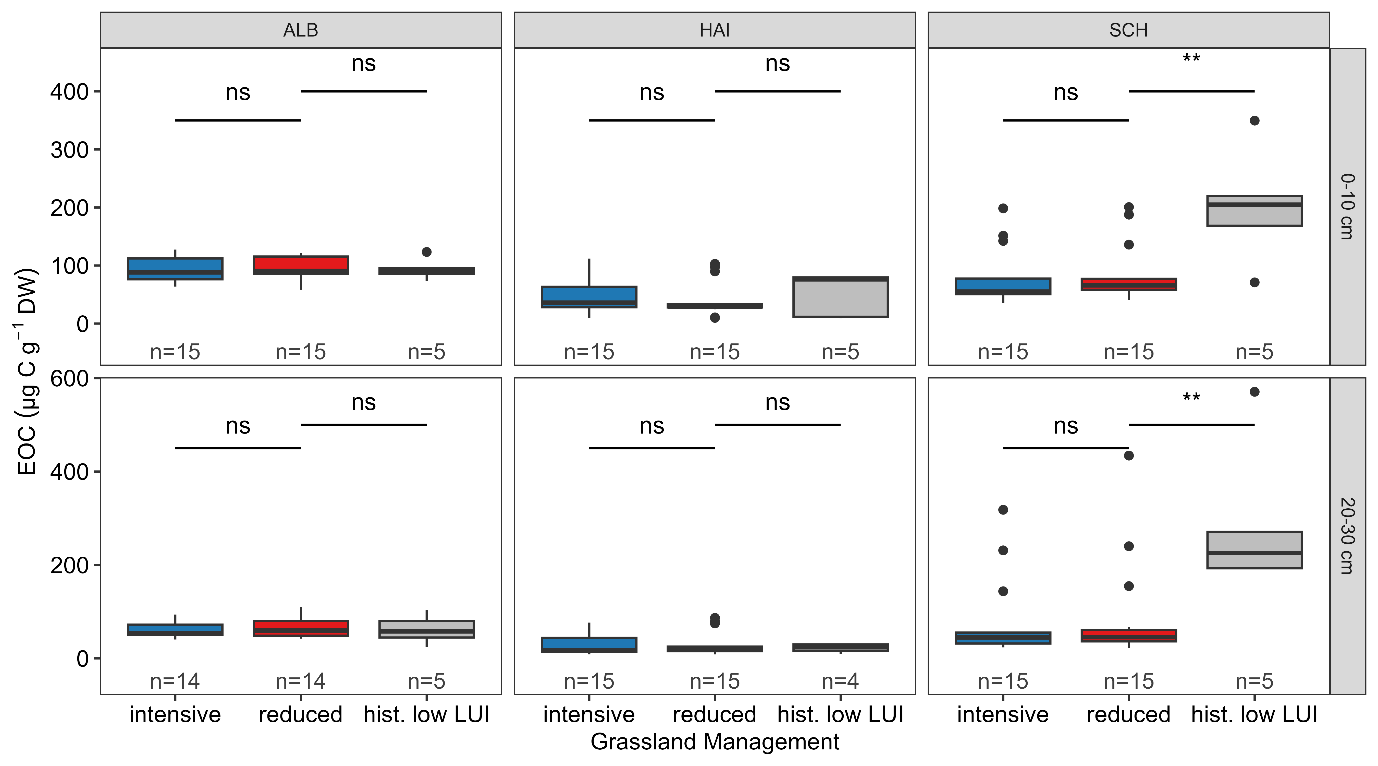

Figure 13: Extractable total nitrogen (ETN) in top- and subsoil in in the regions of Schwäbische Alb (ALB), Hainich (HAI) and Schorfheide (SCH).
